# Differential cell-ECM interaction of rhabdomyosarcoma subtypes regulated by PAX3-FOXO1

**DOI:** 10.1101/2024.06.11.598505

**Authors:** Antonios Chronopoulos, Ivan Chavez, Chandra Vemula, Nikhil Mittal, Vic Zamloot, Sangyoon J Han, JinSeok Park

## Abstract

Rhabdomyosarcoma (RMS) is the most common childhood soft tissue sarcoma, with two subtypes: Fusion-positive RMS (FPRMS), which has the PAX3-FOXO1 fusion gene, and fusion-negative RMS (FNRMS). Despite their distinct characteristics, treatments mainly rely on conventional chemotherapies without considering these differences. This study highlights that FNRMS cells exhibit significantly heightened interaction with the extracellular matrix (ECM) compared to FPRMS cells.

Using single-cell RNA sequencing of skeletal muscle tissues and RNA sequencing of RMS samples, we identified the upregulation of genes related to cell-ECM interaction and TGFβ signaling in FNRMS compared to FPRMS. We also confirmed enhanced cell-ECM interaction stimulated by TGFβ signaling in FNRMS cells, using confocal reflection microscopy to monitor dynamic cell-ECM interaction and a live-cell sensor to quantitatively assess TGFβ signaling activity. Additionally, we discovered that the PAX3-FOXO1 fusion gene, characteristic of FPRMS, stimulated nitric oxide synthesis, which suppresses TGFβ signaling and reduces cell-ECM interaction.

These findings suggest that PAX3-FOXO1 determines the diminished cell-ECM interactions in FPRMS. Experimental data show higher sensitivity of FNRMS to cell-ECM interaction disruption and TGFβ inhibition. Furthermore, the diminished cell-ECM interaction in FPRMS, allowing cells to survive in the ectopic environment through circulation, may partly explain its higher metastatic potential compared to FNRMS.

## Introduction

Rhabdomyosarcoma (RMS) is the most common soft tissue sarcoma in childhood and shares histopathological features with developing skeletal muscle. RMS consists of two major subtypes: fusion-positive RMS (FPRMS), characterized by a chromosomal translocation that results in the oncogenic PAX3-FOXO1 fusion gene, and fusion-negative RMS (FNRMS), which is primarily driven by activating mutations in RTK/RAS pathways ^1^. Despite distinct molecular differences and clinical outcomes between FPRMS and FNRMS, the current treatment protocols do not differentiate between the two subtypes. This is largely due to the lack of understanding of their differential molecular features, which could potentially serve as vulnerable nodes for targeted therapies ^2, 3^. This underscores a need for a more refined understanding of the molecular vulnerabilities underlying RMS subtypes to develop subtype-specific targeted therapeutics.

RMS may arise due to abnormalities in developmental pathways that regulate skeletal muscle determination^2^. RMS cells typically originate from mesenchymal progenitors committed to the myogenic lineage, undergoing inappropriate differentiation arrest at specific developmental stages and leading to persistent proliferation^4^. These findings suggest that FNRMS and FPRMS may have distinct signaling pathways determining differentiation arrests at different stages. Furthermore, the aberrant transcription factor PAX3-FOXO1 may reshape the chromatin landscape and hijack signaling pathways associated with inappropriate differentiation arrest and oncogenic transformation.

The extracellular matrix (ECM), which provides a physical scaffold, plays a crucial role in regulating cell survival and growth through anchorage-dependent signaling pathways^5–7^. In normal myogenic cells, cell-ECM interaction ensures that cells remain anchored to their niche, receiving signals that drive myogenic differentiation ^8–12^. The engagement of ECM in myogenic differentiation suggests its potential role in defining distinct features between FNRMS and FPRMS. Furthermore, in metastasis, ECM-mediated anchorage dependence may prevent tumor dissemination and diminish survival capacity of intravasated circulating tumor cells, which affect metastatic potentials^13^. It remains unclear if and how PAX3-FOXO1 in FPRMS alters transcriptional and signaling networks to reprogram cell-ECM interaction, potentially defining differential signaling profiles between FPRMS and FNRMS. Understanding the molecular mechanisms by which the fusion status of RMS may reprogram anchorage dependence is critical for discerning their different features associated with patients’ outcomes and developing targeted therapies for each RMS subtype.

Herein, we leverage single-cell RNA sequencing (sc-RNA seq) to demonstrate that FNRMS and FPRMS signatures emerge in different cell populations at distinct stages in the overlapping trajectory of normal human skeletal myogenesis. Using a multi-modal approach that combines transcriptomic and functional assays, we found differential cell-ECM interactions between FNRMS and FPRMS. We also demonstrate that PAX3-FOXO1 disrupts cell-ECM interaction to promote anchorage independence by suppressing TGFβ signaling through increased NOS1-mediated synthesis of nitric oxide (NO). We reveal that targeting cell-matrix interaction by disrupting focal adhesions such as FAK and Src or TGFβ signaling could be sufficient to induce growth inhibition in the anchorage-dependent FNRMS subtype. Knockdown of PAX3-FOXO1 in FPRMS, reprogramming anchorage dependency, re-sensitized FPRMS to the growth inhibitory effect of suppressing cell-ECM interaction. We also found that PAX3-FOXO1 may promote an “adhesion-to-suspension transition^13^,” enhancing the survival capacity of circulating tumor cells and increasing metastatic potential, which may, at least partially, explain the higher metastatic propensity of FPRMS compared to FNRMS.

## Result

### FPRMS and FNRMS-like subtypes are enriched in distinct stages of muscle development from embryo to adult

Wei *et al.* demonstrated that the distinct developmental stages at which myogenic differentiation becomes arrested exhibit RMS-subtype specific transcriptional programs^4^. We hypothesized that these distinct arrested stages determine divergent characteristics, including underlying signaling pathways and cell phenotypes, between FNRMS and FPRMS. To examine this hypothesis, we re-analyzed scRNA-seq data of limb muscle tissues obtained from embryonic (6-8 weeks), fetal (9-18 weeks), and juvenile/adult stages (7-42 years) (GSE147457) (**Figure 1a** and **Supplementary Figure 1a**).^14^ The Leiden clustering of the scRNA-seq data^15^ identified ten distinct subgroups based on the similarities in gene expression (**Figure 1b** and **Supplementary Table 1**) and differentially expressed genes (DEGs) of each subgroup (**Supplementary Table 2**).

**Figure 1.**
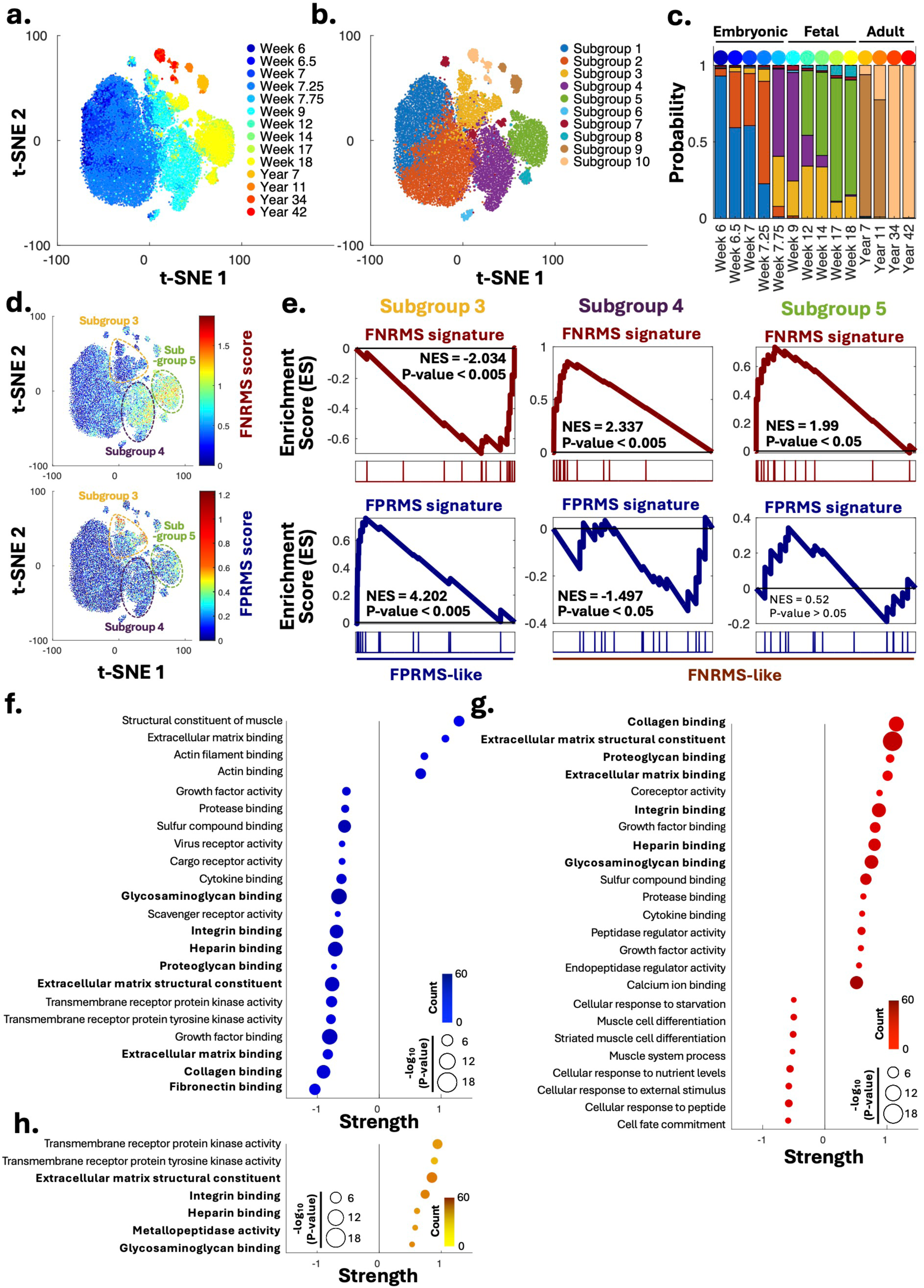
FPRMS and FNRMS mirror different subgroups in myogenic development. **(a)** t-distributed Stochastic Neighboring Embedding (t-SNE) rendering single-cell RNA (scRNA) sequencing of skeletal muscle tissues obtained from human prenatal development (from 6 to 18 weeks) and juveniles/adults (from 7 to 42 years). scRNA sequencing data is from GSE147457. **(b)** Clusters identified by the Leiden algorithm, representing subgroups in myogenic development. **(c)** Population probability of cells in each subgroup of each week’s tissue sample. **(d)** Expression of FNRMS signatures (FNRMS score, top) and FPRMS signatures (FPRMS score, bottom) in the subgroups. **(e)** Gene Set Enrichment Analysis (GSEA) of highly expressed genes in Subgroup 3, 4, and 5 with FNRMS and FPRMS signatures as references, suggesting molecular similarity of Subgroup 3 with FPRMS and the similarity of Subgroup 4 and 5 with FNRMS. A positive enrichment score (ES) and normalized ES indicate the similarity of inquired genes with the reference gene set. Gene Ontology (GO) analysis of **(f)** significantly low expressed genes in Subgroup 3, **(g)** highly expressed genes in Subgroup 4, and **(h)** highly expressed genes in FNRMS compared to FPRMS, using GO Function Gene sets as the reference gene set. Color represents the count of inquired genes overlapping with each specific reference gene set. Circle size indicates adjusted P-value, and strength means the ratio of inquired to reference gene counts, implying the enhanced expression of the corresponding reference. Gene expression data of FPRMS and FNRMS is from GSE114621.

We mapped the distribution enrichment of the subgroups to the developmental trajectory from embryo to adult (**Figure 1c**). Subgroups 1 to 5, substantially spanning from embryonic and fetal status, share proximal connections within the trajectory (**Supplementary Figure 1b**). We discovered that Subgroups 1 and 2, enriched in early embryonic stages (Week 6-7.25), demonstrated higher expression of genes indicative of early development, e.g., embryo (Subgroup 1) and heart/head/axon guidance (Subgroup 2) (**Supplementary Table 3**), through Gene Ontology (GO) analysis with the DEGs. Subgroups 3 to 5, which may represent later embryonic or fetus stages relative Subgroups 1 and 2, exhibit molecular features indicating more differentiated tissues, muscle skeletal system (Subgroup 3), fetus and cardiovascular system (Subgroup 4 and 5) (**Supplementary Table 3**). Additionally, we assess the Fetal score, the expression levels of highly expressed genes in Subgroup 5, highly enriched in fetal tissues, and the Embryonic score, those of highly expressed genes in Subgroup 1, highly enriched embryonic tissues, of each subgroup. Increased Fetal scores and reduced Embryonic scores of Subgroups 3 to 5 compared to Subgroups 1 and 2 suggest that they represent later developmental stages (**Supplementary Figure 1c-e**).

Next, we assessed the enrichment of FN and FPRMS core signatures, defined by Gryder *et al.*^16^ (**Supplementary Table 4**), in the identified subgroups (**Figure 1d**). Interestingly, we discovered that the FPRMS core signature was enriched in Subgroup 3, while the FNRMS core signature was enriched primarily in Subgroups 4 and 5. Gene set enrichment analysis (GSEA) shows robust upregulation of FPRMS signatures in Subgroup 3 with a concurrent depletion of FNRMS signatures (**Figure 1e** and **Supplementary Figure 1f**). A reverse trend is observed for Subgroup 4, showing depletion of FPRMS signatures and upregulation of FNRMS signatures. Subgroup 5 shows a more modest upregulation in FNRMS signatures compared to Subgroup 4 and, similarly, a less pronounced depletion of FPRMS signatures. Wei *et al.* discovered the enrichment of FPRMS core signatures at the stage that embryonic muscle transits to fetal muscle at Week 7-7.5^4^. Our analysis consistently exhibited that FPRMS-like Subgroup 3, expressing skeletal muscle-related genes, substantially started to emerge around Week 7.5 (**Supplementary Table 3** and **Figure 1c**). Furthermore, FNRMS-like Subgroups 4 and 5 exhibited the expression of genes defining vascular system (**Supplementary Table 3**), in line with previous reports indicating an endothelial cell as an origin of cells for FNRMS^17^. Conversely, higher expression of genes indicating skeletal muscle in Subgroup 3 may reflect myogenic reprogramming driven by the PAX3-FOXO1 fusion gene of FPRMS^18^. These results support the idea that the development of both RMS subtypes aligns with muscle development and each subtype employs specific transcriptional programs at its different stages To test whether the distinct molecular features of the most FPMRS-like Subgroup 3 and the most FNRMS-like Subgroup 4 mirror underlying differential features between RMS subtypes, we investigated DEGs in the subgroups (**Supplementary Table 2** and **3**) and each RMS subtype (**Supplementary Figure 1f** and **Supplementary Table 5** and **6**) followed by GO analysis. Intriguingly, functional annotation of inquired DEGs in Subgroups 3 and 4 revealed genes involved in cell-extracellular matrix (ECM) interactions to be strongly downregulated for Subgroup 3 (**Figure 1f**). By contrast, cell-ECM interaction gene sets were strongly upregulated in Subgroup 4 (**Figure 1g**). Head-to-head comparison between the two RMS subgroups shows gene sets involved in cell-ECM interactions (e.g., ECM constituents, integrin binding, heparin binding) to be enriched in FNRMS relative to FP-RMS (**Figure 1h**), consistent with the similarity of the subgroups corresponding to each RMS subtype. Thus, we hypothesize that FNRMS cells have higher cell-ECM interaction than FPRMS cells.

### FNRMS shows higher cell-ECM interaction than FPRMS

To evaluate our hypothesis of higher cell-ECM interaction in FNRMS than in FPRMS, we performed immunofluorescent staining for the F-actin cytoskeleton and phosphorylated focal adhesion kinase (P-FAK), indicating focal adhesion formations in FNRMS (RD and SMS-CTR) and FPRMS (Rh30 and Rh41) cell lines. We found that FNRMS exhibits prominent stress fibers necessary for establishing and maintaining cell-ECM interactions, as opposed to FPRMS, which is largely devoid of cytoskeletal stress fiber bundles. FNRMS also showed elevated focal adhesion formations indicated by P-FAK, representing enhanced cell-ECM interaction (**Figure 2a**). These differences could arise from cell-intrinsic differences between RMS subtypes in signaling pathways regulating cytoskeletal organization and focal adhesion, which determine cell-ECM interaction and, accordingly, related cell phenotypes ^19–21^. Indeed, the higher cell-ECM interaction of FNRMS than FPRMS was further corroborated by an increased cell spreading area in FNRMS relative to FPRMS (**Figure 2b**).

**Figure 2.**
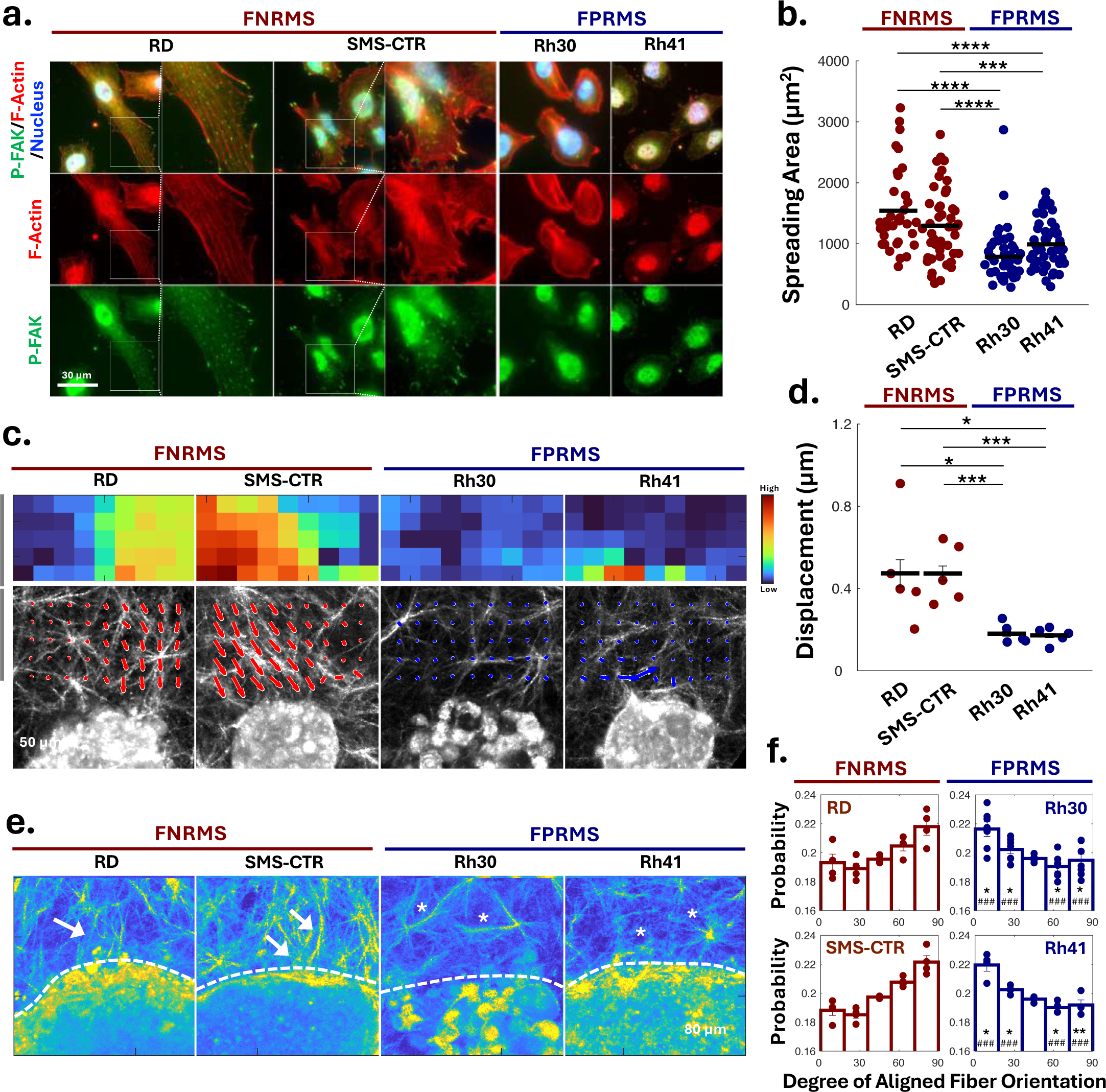
FNRMS shows higher cell-ECM interaction than FPRMS. **(a)** Immunofluorescent images of FNRMS (RD and SMS-CTR) and FPRMS (Rh30 and Rh41) cell lines, stained for F-actin (with phalloidin) and phospho-focal adhesion kinase (P-FAK), displaying focal adhesion. **(b)** Cell spreading areas for each RMS subtype cell line. Each dot represents the value of a corresponding cell and lines are averages of each cell line. Error bars represent S.E. Statistical significance was assessed using two-sided Student’s t-test: ***P<0.005, ****P<0.001. **(c)** Initial 3-hour collagen fiber displacement near spheroids, monitored by confocal reflection microscopy (CRM) and analyzed by particle image velocimetry (PIV). Top: pseudo-colored images showing the displacement degree of collagen fibers caused by spheroids. Bottom: Arrows indicate the direction and magnitude of collagen fiber displacement. **(d)** Quantification of the collagen fiber displacements. Error bars represent S.E. Statistical significance was assessed using two-sided Student’s t-test: *P<0.05 and ***P<0.005. **(e)** Collagen fiber alignment 16 hours post-gelation with spheroids. Dashed lines denote spheroid edges. Arrows point to the fibers aligned perpendicularly to FNRMS spheroid edges, while asterisks highlight the fibers aligned parallel to FPRMS spheroid edges. **(f)** Distribution of collagen fiber orientation around RMS cell spheroids. Angles at 90° represent perpendicular alignment to spheroid edges; 0° represents parallel alignment. Dots represent individual spheroid values. Asterisks (*) denote statistical differences from RD spheroid values and pounds (#) indicate differences from SMS-CTR spheroid values, assessed using two-sided Student’s t-test: *P<0.05, **P<0.01, ^###^P<0.005. ****P<0.001.

To verify the differential cell-ECM interaction more functionally, we utilized confocal reflection microscopy (CRM), a novel microscopic technique to characterize the spatial and temporal dynamics of cell-ECM interactions within 3D collagen matrix in three-dimensional (3D) culture^22, 23^. Using hanging drop method^24^, we generated tumor spheroids of FPRMS and FNRMS cell lines and embedded them in the 3D gels of collagen type, the representative ECM, before initiating time-lapse CRM imaging. We quantified collagen fiber displacement and alignment as metrics to assess the extent to which the spheroids physically interact with and remodel the collagen matrix through particle image velocimetry (PIV)^25, 26^. We found collagen fibers near the FNRMS spheroids exhibit a higher displacement magnitude compared to those near FPRMS spheroids (**Figure 2c,d** and **Supplementary Movie 1**). We next assessed collagen fiber alignment via matrix remodeling at the boundary edge of the tumor spheroids. Fiber orientation distribution analysis showed a higher proportion of perpendicularly aligned collagen fibers (relative to the edge) of FNRMS spheroids compared to FPRMS (**Figure 2e, f, Supplementary Figure 2a**, and **b**). The perpendicular alignment of ECM collagen fibers implies that FNRMS cells exert pulling forces reorganizing ECM topographically, indicating strong cell-ECM interaction^27^. In contrast, their parallel alignment to the edge of FPRMS spheroids indicates passive radial expansion of the spheroid largely devoid of active pulling forces by the spheroid growth and no active cell-ECM interaction. Additionally, we measured the traction forces of SMS-CTR (FNRMS) and Rh41 (FPRMS) cells using traction force microscopy (**Supplementary Figure 2c** and **d**)^28^. Considering the traction force of cells is indicative of cell-ECM interaction^29, 30^, the higher traction force of FNRMS than FPRMS aligns well with their higher cell-ECM interaction. Collectively, these results indicate FNRMS exhibits stronger cell-ECM interaction relative to FPRMS.

### TGFβ signaling, highly activated in FNRMS, stimulates cell-ECM interaction

To identify candidate pathways responsible for the differential cell-ECM interaction between the two RMS subtypes, we conducted GO analysis of the DEGs in Subgroups 3 and 4, using the KEGG database as a reference data set. Interestingly, the TGFβ signaling pathway was downregulated in FPRMS-like Subgroup 3 and upregulated in FNRMS-like Subgroup 4 (FNRMS) (**Figure 3a**). TGFβ regulates ECM synthesis and remodeling, promoting cell-ECM interaction^31, 32^. The activation of TGFβ signaling involves the canonical pathway where TGFβ binds to its cognate receptor on the cell surface, leading to the activation of downstream SMAD proteins. These SMAD proteins, including Smad3, translocate into the nucleus and regulate the transcription of target genes involved in cytoskeletal organization and ECM remodeling^33, 34^. A previous report demonstrated that ERMS (corresponding to FNRMS) displayed a higher response to TGFβ than ARMS (corresponding to FPRMS), implicating higher TGFβ signaling activity in FNRMS^35^. Concomitantly, DEGs in FNRMS exhibited higher enrichment of genes associated with TGFβ signaling (**Figure 3a**). Thus, we hypothesized that FNRMS has higher activity of TGFβ signaling relative to FPRMS, resulting in its heightened cell-ECM interaction.

**Figure 3.**
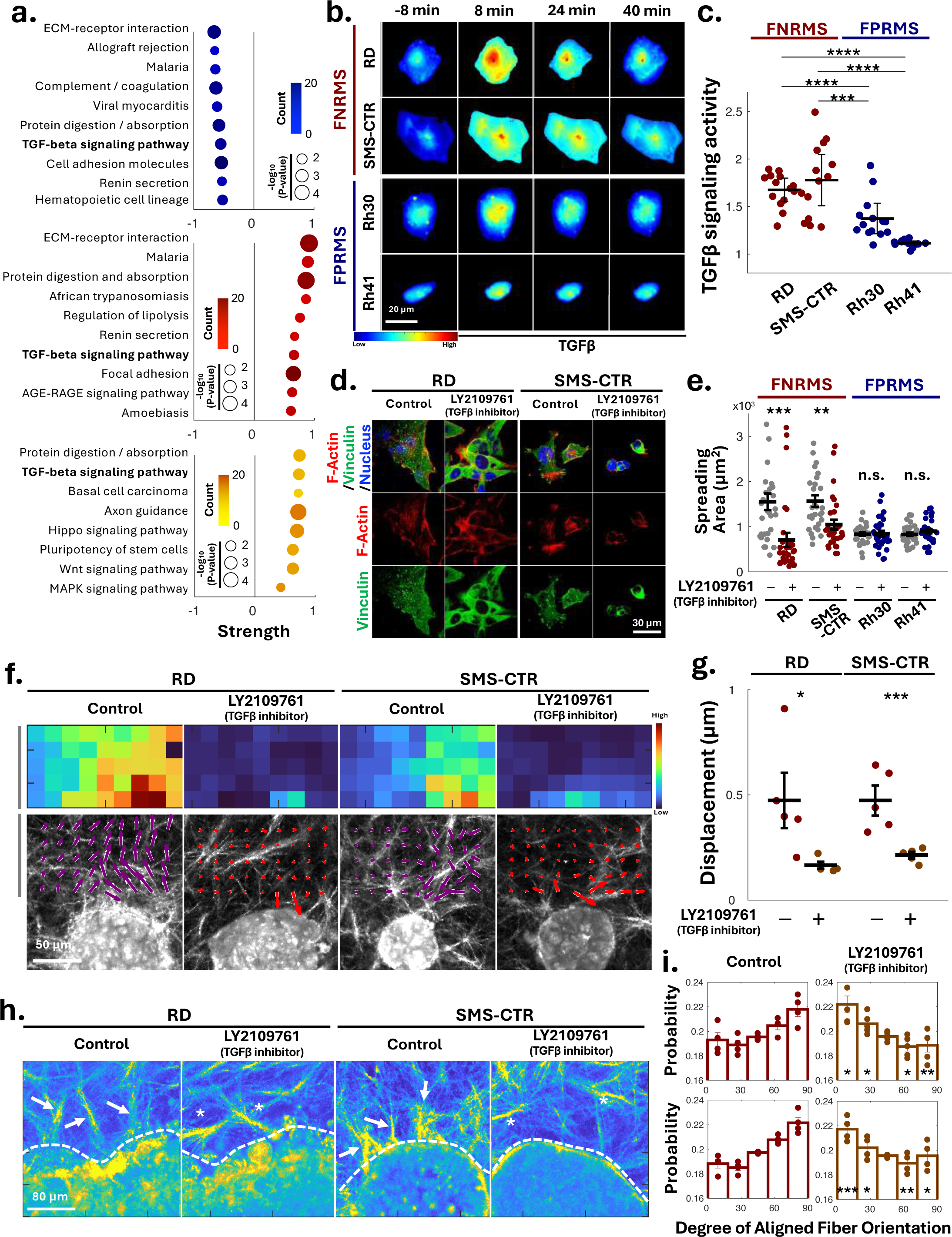
FNRMS exhibits higher TGFβ signaling activity than FPRMS. **(a)** GO analysis of genes significantly low expressed genes in Subgroup 3, reflecting FPRMS (top), high-expressed genes in Subgroup 4, reflecting FNRMS (middle), and genes highly expressed in FNRMS compared to FPRMS (bottom), using KEGG Gene Sets as the reference gene set. We highlight TGFβ signaling as a suppressed signaling pathway in Subgroup 3, an activated signaling pathway in Subgroup 4, and an enhanced signaling pathway in FNRMS compared to FPRMS. Color represents the count of inquired genes overlapping with each specific reference gene set. Circle size indicates P-value, and strength means the ratio of inquired to reference gene counts, implying the enhanced expression of the corresponding reference. **(b)** Dynamic Smad3 nuclear localization upon TGFβ treatment (4μg/ml) in FNRMS and FPRMS cell lines, monitored by using NG-Smad3 live cell sensor. **(c)** TGFβ signaling activity, quantified by the ratio of intensity in nuclear Smad3 after TGFβ treatment to before the treatment, of FNRMS and FPRMS cell lines. Each dot represents the value of a corresponding cell and lines are averages of each cell line. Error bars represent S.E. Statistical significance was assessed using two-sided Student’s t-test: ***P<0.005, ****P<0.001. **(d)** Immunofluorescent images of RD and SMS-CTR (FNRMS), stained for F-actin (with phalloidin) and vinculin, showing focal adhesion, with and without the treatment of a TGFβ inhibitor (LY2109761, 100nM, 12 hours). **(e)** Cell spreading areas for FNRMS and FPRMS cell lines after the treatment with the TGFβ inhibitor. Each dot represents the value of a corresponding cell and lines are averages of each cell line. Error bars represent S.E. Statistical significance was assessed using two-sided Student’s t-test: **P<0.01, ***P<0.005, and n.s. = no significance. **(f)** Collagen fiber displacement near FNRMS spheroids with and without the treatment of LY2109761 (100 nM), monitored by CRM and analyzed by PIV. Top: pseudo-colored images showing the displacement degree of collagen fibers caused by FNRMS spheroids. Bottom: Arrows indicate the direction and magnitude of collagen fiber displacement. **(g)** Quantification of the collagen fiber displacements surrounding FNRMS spheroids with and without TGFβ inhibition. Error bars represent S.E. Statistical significance between control and LY21097861 was assessed using two-sided Student’s t-test: *P<0.05 and ***P<0.005. **(h)** Collagen fiber alignment 16 hours post-gelation with FNRMS spheroids with and without the treatment of LY2109761. Dashed lines denote FNRMS spheroid edges. Arrows point to the fibers aligned perpendicularly to the spheroid edges, while asterisks highlight the fibers aligned parallel to the spheroid edges. **(i)** Distribution of collagen fiber orientation around FNRMS spheroids with and without the TGFβ inhibitor. Dots represent individual spheroid values. Asterisks (*) denote statistical differences of values between spheroids with and without LY2109761, assessed using two-sided Student’s t-test: *P<0.05, **P<0.01, **P <0.005.

To confirm our hypothesis, we employed live cell imaging to monitor the relative change in nuclear SMAD3 signal, indicative of TGFβ signaling activation, upon stimulation with recombinant TGFβ ligand (4 ng/mL), using the NeonGreen(NG)-Smad3 live-cell sensor^36^ (**Figure 3b**). Smad3 expression varied across cells, limiting our ability to assess cell-to-cell TGFβ signaling activity ^37^. However, the relative change of the live-cell sensor, representing TGFβ signaling activity robustly, allows us to overcome its variation across various cell lines. Intriguingly, the NG-Smad3 dynamics revealed a marked increase in TGFβ responsiveness across both FNRMS cell lines. In contrast, the FPRMS cell lines responded weakly to ligand stimulation compared to FNRMS cells (**Figure 3c** and **Supplementary Figure 3**). To evaluate if increased TGFβ signaling activity in FNRMS affects cell-ECM interaction, we treated FNRMS cell lines with the selective TGFβ receptor type I/II (TβRI/II) dual inhibitor LY2109761, followed by immunofluorescent staining for F-actin and vinculin at focal adhesions. Inhibition of TβRI/II disrupted stress fibers, characterized by a predominant localization of actin at the cell cortex. It also led to a marked decrease in the puncta of focal adhesion, indicated by vinculin (**Figure 3d**). TβRI/II inhibition reduced the cell spreading area in FNRMS but not FPRMS, supporting our hypothesis that highly-activated TGFβ signaling in FNRMS enhances its cell-ECM interaction. LY2109761-treated FNRMS spheroids also exhibited a diminished capacity to physically interact with the surrounding collagen matrix, as indicated by reduced displacement and alignment of collagen fibers observed through CRM (**Figure 3f-I** and **Supplementary Movie 2**). These results suggest that TGFβ inhibition associated with the FNRMS subtype causes cytoskeletal and adhesive changes resulting in reduced cell-ECM interactions.

### PAX3-FOXO1 of FPRMS suppressed TGFβ signaling and cell-ECM interaction

The *PAX3-FOXO1* fusion gene is a critical molecular signature in FPRMS, differentiating it from FNRMS. We hypothesized that the fusion gene suppresses TGFβ signaling and cell-ECM interaction. To evaluate this hypothesis, we generated stable FPRMS cell lines expressing a doxycycline-inducible lentivirus expressing shRNA directed against PAX3-FOXO1 breakpoint (PF^KD^)^38^ (**Supplementary Figure 4a**). We identified DEGs highly in PF^KD^ cells compared to non-induced control cells (**Supplementary Table 7**). Interestingly, we discovered that the DEGs are highly expressed in FNRMS-like Subgroups 4 and 5, relative to FPRMS-like Subgroup 3 identified through scRNA-seq from muscle biopsies across development. (**Figure 4a** and **Supplementary Figure 4b**). The DEGs consistently demonstrated positive enrichment in FNRMS signatures, although there was no statistically negative significant enrichment in FPRMS signatures (**Supplementary Figure 4c**). GSEA analysis also confirmed a distinct enrichment of upregulated genes in PF^KD^ cells within the Subgroup 4 (**Figure 4b**). Furthermore, GO analysis of DEGs in PF^KD^ revealed an upregulation of genes involved in cell-ECM interactions and cell adhesion (**Figure 4c** and **Supplementary Table 8**).

**Figure 4.**
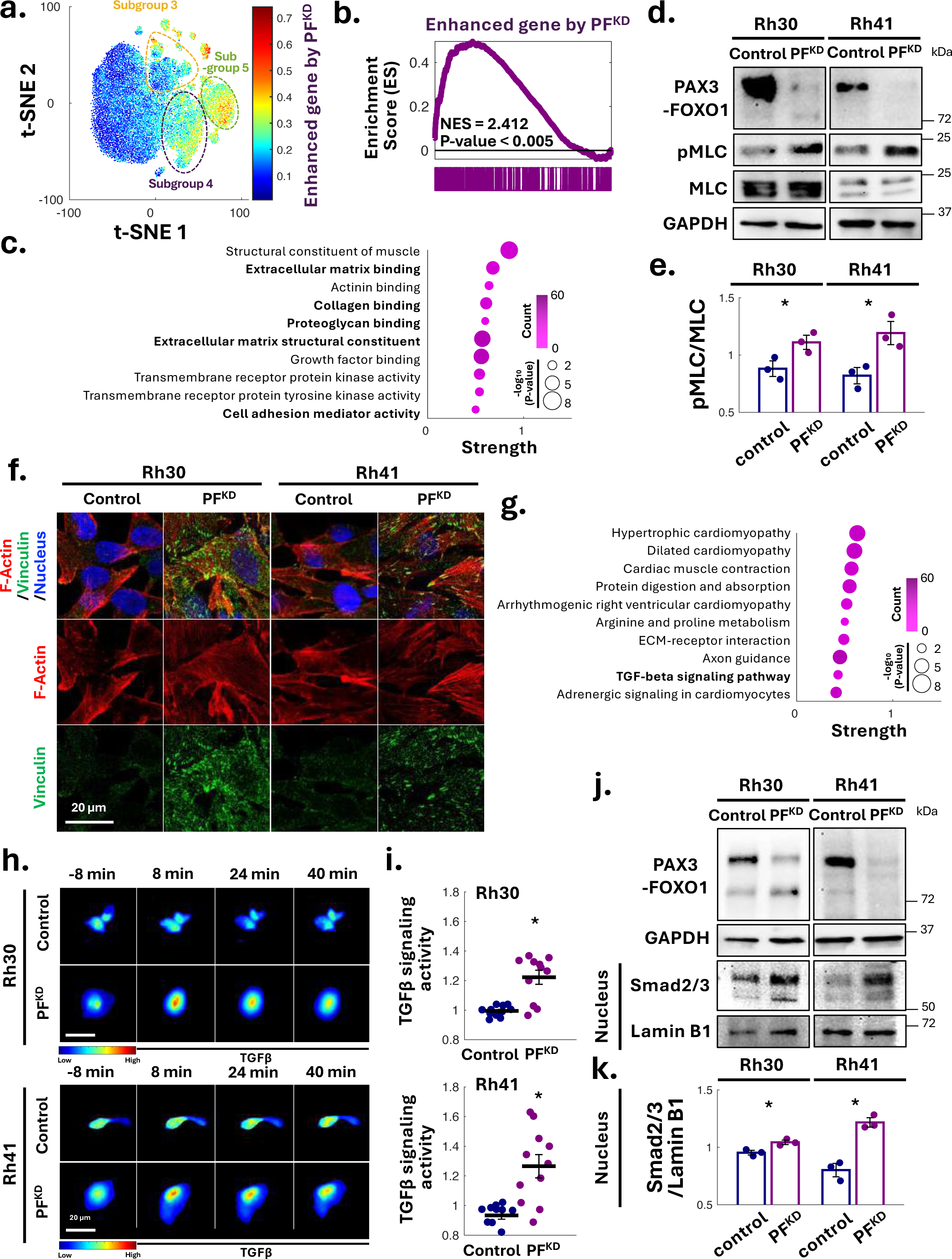
PAX3-FOXO1 suppresses TGFβ signaling and cell-ECM interaction. **(a)** Higher expression of genes enhanced by PAX3-FOXO1 knockdown (PF^KD^) in FPRMS cells and in FNRMS-like Subgroup 4 and 5, than in FPRMS-like Subgroup 3. The color indicates the average expression level of genes highly expressed in PF^KD^ FPRMS cells compared to control cells in each cell. **(b)** GSEA suggests that highly expressed genes in Subgroup 4 exhibit statistically significant overlap with genes enhanced by PF^KD^ in FPRMS cells. **(c)** GO analysis of highly expressed genes in PF^KD^ FPRMS cells (Rh30 and Rh41) compared to control cells, using GO Function Gene sets as the reference gene set. **(d)** Immunoblotting results and **(e)** its quantification shows higher phosphorylation of myosin light chain (pMLC) of PFKD FPRMS cells than control cells, implying their higher cell-ECM interaction. Error bars represent S.E. Statistical significance was assessed using two-sided Student’s t-tests: *P<0.05 (n = 3). **(f)** Immunofluorescent images of PF^KD^ Rh30 and Rh41, stained for F-actin (with phalloidin) and vinculin, showing focal adhesion. **(g)** GO analysis of highly expressed genes in PF^KD^ FPRMS cells (Rh30 and Rh41) compared to control cells, using KEGG Gene sets as the reference gene set. In **(c)** and **(g)**, color represents the count of inquired genes overlapping with each specific reference gene set. Circle size indicates adjusted P-value, and strength means the ratio of inquired to reference gene counts, implying the enhanced expression of the corresponding reference. **(h)** Dynamic Smad3 nuclear localization upon TGFβ treatment (4µg/ml) in PF^KD^ FPRMS cell lines, monitored using NG-Smad3 live-cell sensor. **(i)** TGFβ signaling activity, quantified by the ratio of intensity in nuclear Smad3 after TGFβ treatment to before the treatment, in PF^KD^ FPRMS cells. Each dot represents the value of a cell and lines are averages of each cell line. Error bars represent S.E. Statistical significance was assessed using two-sided Student’s t-tests: *P<0.05. **(j)** Immunoblotting of nuclear fractionized proteins and **(k)** its quantification exhibit higher Smad3 nuclear localization of PF^KD^ FPRMS cells than control cells, implying their higher TGFβ signaling activity. Error bars represent S.E. Statistical significance was assessed using two-sided Student’s t-tests: *P<0.05 (n = 3).

Next, we examined that PF^KD^ FPRMS cells demonstrated the features of enhanced cell-ECM interaction as suggested by sequencing analysis. PF^KD^ cells had significantly increased phosphorylation of myosin light chain (MLC), establishing cytoskeletal stress fibers ^39^ and maturing focal adhesion^40^, which represents higher cell-ECM interaction (**Figure 4e**). Furthermore, PF^KD^ cells had a notably increased number of vinculin-enriched focal adhesions and stress fiber bundles relative to the scramble counterparts, indicating cytoskeletal remodeling and stronger cell-matrix interactions (**Figure 4f** and **Supplementary Figure 4d**). GO analysis on DEGs in PF^KD^ cells showed significant upregulation of the TGFβ pathway using the GO Function gene sets as a reference set (**Figure 4g** and **Supplementary Table 8**). Using the NG-Smad3 live-cell sensor, we noted increased TGFβ responsiveness for both PF^KD^ Rh30 and Rh41 cell lines relative to controls (**Figure 4h, i**, **Supplementary Figure 5a** and **b**). Immunoblotting of the nuclear fraction also confirmed elevated Smad2/3 protein expression in PF^KD^ FPRMS cell lines, implicating higher nuclear localization of the SMAD proteins, indicative of higher TGFβ signaling activation (**Figure 4j, k**, and **Supplementary Figure 5c**).

PF^KD^ cells exhibited increased spreading area relative to their control counterparts (**Figure 5a** and **Supplementary Figure 6a**). However, upon pharmacological inhibition of TβRI/II in PF^KD^ cells, this effect was nullified. In line with these findings, imaging using CRM of PF^KD^ spheroids embedded in collagen revealed increased interaction between cells and the ECM, as indicated by greater displacement of collagen fibers compared to control counterparts (**Figure 5b, c, Supplementary Figure 6b**, and **Supplementary Movie 3**). However, this effect was also nullified upon pharmacological inhibition of TβRI/II with LY2109761. The same trend was observed for cell-mediated alignment of collagen fibers, where PF^KD^ possessed a greater ability to re-organize the matrix by aligning the surrounding fibers relative to their scramble controls, and this effect was similarly abolished with LY2109761 treatment (**Figure 5d, e**, and **Supplementary Figure 6c**). These results suggest that the PAX3-FOXO1 fusion gene disrupts cell-ECM interactions by inhibiting TGFβ signaling.

**Figure 5.**
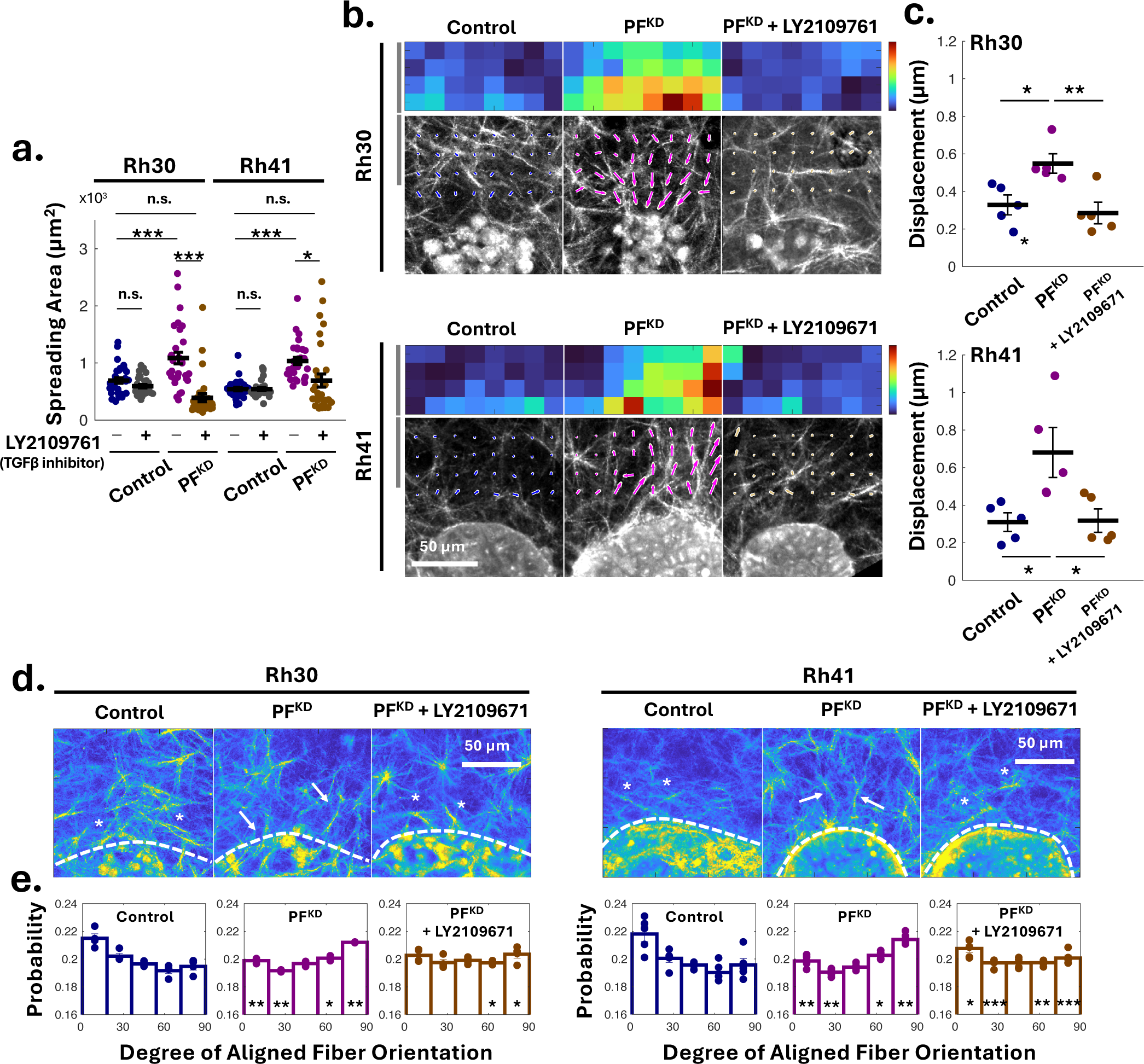
The knockdown of PAX3-FOXO1 enhances cell-ECM interaction via TGFβ signaling. **(a)** Cell spreading areas of PF^KD^ FPRMS cell lines upon the treatment of the TGFβ inhibitor, LY2109761 (100 nM, 12 hours). **(b)** Collagen fiber displacement near spheroids of control FPRMS cells (left), PF^KD^ FPRMS cells (middle), and PF^KD^ FPRMS cells with the treatment of LY2109761 (100 nM, right), monitored by CRM and analyzed by PIV. Top: pseudo-colored images showing the displacement degree of collagen fibers caused by spheroids. Bottom: Arrows indicate the direction and magnitude of collagen fiber displacement. **(c)** Quantification of the collagen fiber displacements of PF^KD^ FPRMS cell lines upon treatment of the TGFβ inhibitor. In **(a)** and **(c)**, each dot represents the value of a corresponding cell or spheroid. Lines are averages of each condition. Error bars represent S.E. Statistical significance was assessed using two-sided Student’s t-tests: *P<0.05, **P<0.01, ***P>0.005, and n.s = no significance **(d)** Collagen fiber alignment 16 hours post-gelation with control FPRMS cells (left), PF^KD^ FPRMS cells (middle), and PF^KD^ FPRMS cells with the treatment of LY2109761 (100 nM, right). Dashed lines denote the spheroid edges. Arrows point to the fibers aligned perpendicularly to the spheroid edges, while asterisks (*) highlight the fibers aligned parallel to the edges. **(e)** Distribution of collagen fiber orientation around the spheroids with and without the TGFβ inhibitor. Dots represent individual spheroid values. Statistical differences were assessed using two-sided Student’s t-test: *P<0.05, *P<0.01, and *P<0.005.

### PAX3-FOXO1-mediated nitric oxide synthesis inhibits TGFβ signaling

We then sought to investigate how the PAX3-FOXO1 fusion gene regulates the TGFβ signaling pathway. The fusion gene acts as a rogue transcription factor that can reprogram myogenic development. Gryder *et al.* identified the PAX3-FOXO1 target genes through chromatin immunoprecipitation sequencing (ChIP-seq)^16^. Among the target genes, NOS1 exhibited the most significant positive-fold change in expression in FPRMS compared to FNRMS (**Figure 6a**). In line with this, we also discovered the greatest decrease in the expression of NOS1 in PF^KD^ Rh30 and Rh41 cells compared to controls (**Figure 6b**). NOS1 (Nitric oxide synthases 1) is an enzyme crucial in nitric oxide (NO) synthesis. Interestingly, NO has been considered as disrupting focal adhesion formations^41, 42^ and reducing ECM synthesis^43^ in vascular systems, which remains understudied in cancer research. Furthermore, NO can suppress TGFβ signaling in various context^44–46^. Thus, we hypothesize that NO concentration enhanced by the PAX3-FOXO1 fusion gene expression reduces TGFβ signaling activity and interferes with cell-ECM interaction.

**Figure 6.**
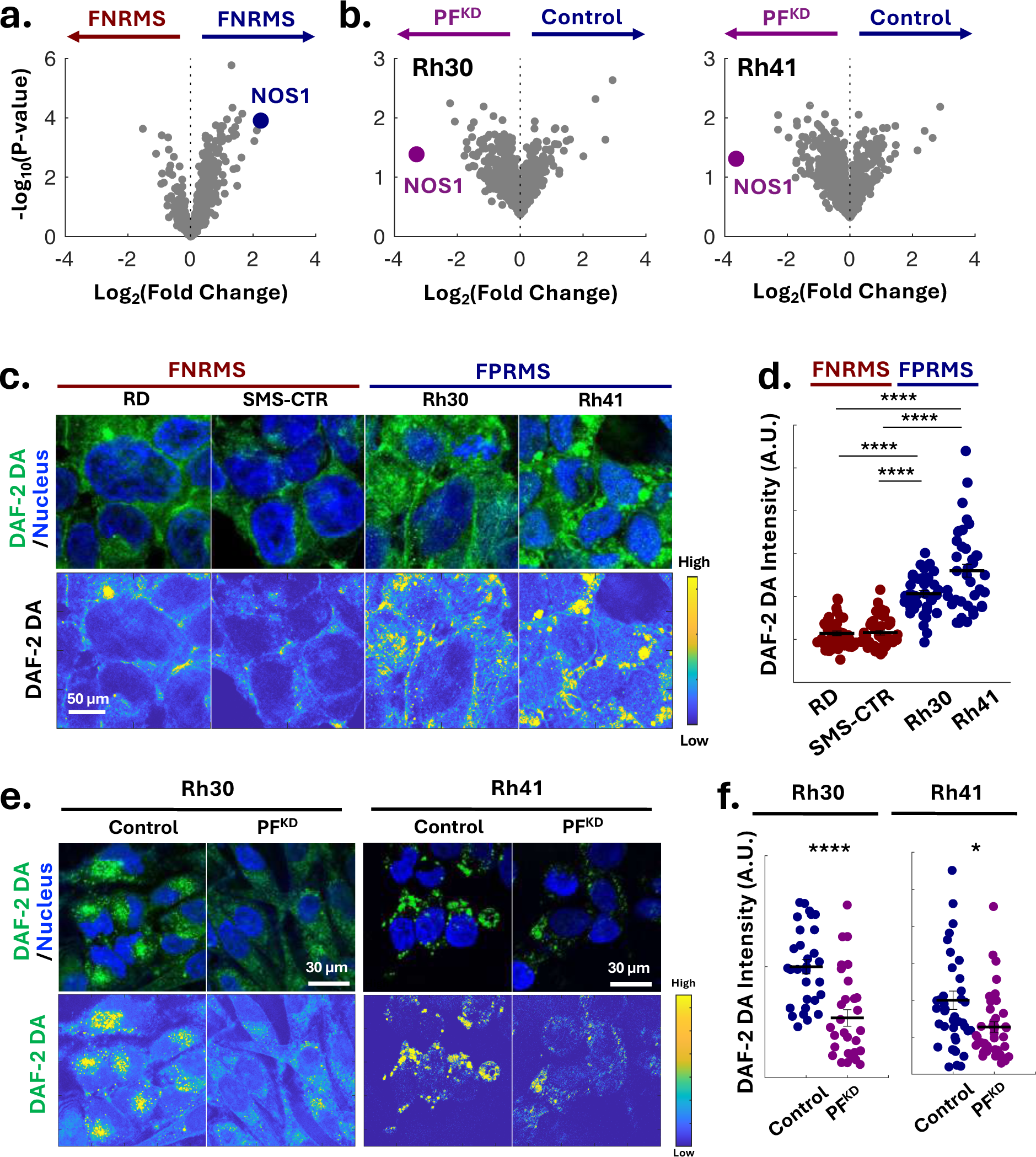
PAX3-FOXO1 stimulates the production of Nitric Oxide (NO) **(a)** Higher NOS1 expression in FPRMS than FNRMS. **(b)** Increased NOS1 expression by PF^KD^. Differentially expressed genes among PAX3-FOXO1 target genes between PF^KD^ Rh30 and control Rh30 (left) and between PF^KD^ Rh41 and control Rh41 (Right). PAX3-FOXO1 target genes are identified in Gryder *et al*. (PMID: 28446439). NOS1 is the gene that commonly exhibits the greatest fold change. **(c)** DAF2-DA staining and its pseudo-color images that probe nitric oxide (NO) in FNRMS and FPRMS cell lines and **(d)** its intensity indicating NO concentration in each cell line, suggesting higher NO concentration in FNRMS than FPRMS cells. **(e)** DAF2-DA staining of PF^KD^ FPRMS cell lines and **(f)** its intensity indicating their NO concentrations, implying higher NO concentration in PF^KD^ FPRMS than control cells. In **(d)** and **(f)**, each dot represents the intensity of a corresponding cell and lines are averages of each cell line. Error bars represent S.E. Statistical significance was assessed using two-sided Student’s t-tests: *P<0.05, and ****P>0.001.

To validate this hypothesis, we first confirmed that both FPRMS cell lines showed increased intracellular NO concentrations relative to FNRMS counterparts, using the fluorescent probe DAF-2 DA that detects the NO levels ^47^ (**Figure 6c, d**). Correspondingly, PF^KD^ cells showed a reduction in the NO levels relative to control FPRMS cells (**Figure 6e, f** and **Supplementary Figure 7**). These results highlight that PAX3-FOXO1 stimulates NOS1-mediated NO synthesis.

To investigate if PAX3-FOXO1-mediated NO synthesis via NOS1 serves as an upstream negative regulator of TGFβ signaling and cell-ECM interaction, we performed immunofluorescent staining for nuclear SMAD3 in Rh41 FPRMS cells with the treatment of NOS1 inhibitor, ARL17477. We demonstrated a marked increase in nuclear SMAD3 intensity upon pharmacological NOS1 inhibition relative to control (**Figure 7a, b)**, further corroborated by immunoblotting showing elevated SMAD2/3 in the nuclear fraction upon the inhibition (**Figure 7c, d)**. These results support our hypothesis, confirming that the NOS1 inhibition stimulates TGFβ signaling. The inhibition also increased F-actin stress fibers and the number and size of vinculin-enriched focal adhesions (**Figure 7e)**. Moreover, NOS1 inhibition led to increased phosphorylation of MLC and cell spreading area relative to untreated FPRMS controls (**Figure 7f-h**). NOS1 inhibition of FPRMS spheroids embedded in collagen-I gels also enhanced their ability to physically interact with the matrix, resulting in higher displacement and alignment of collagen fibers as visualized by CRM (**Figure 7i-k** and **Supplementary Movie 4**). Taken together, these findings imply that PAX3-FOXO1 disrupts cell-ECM interaction by inhibiting TGFβ signaling via the PAX3-FOXO1-stimulated synthesis of NO.

**Figure 7.**
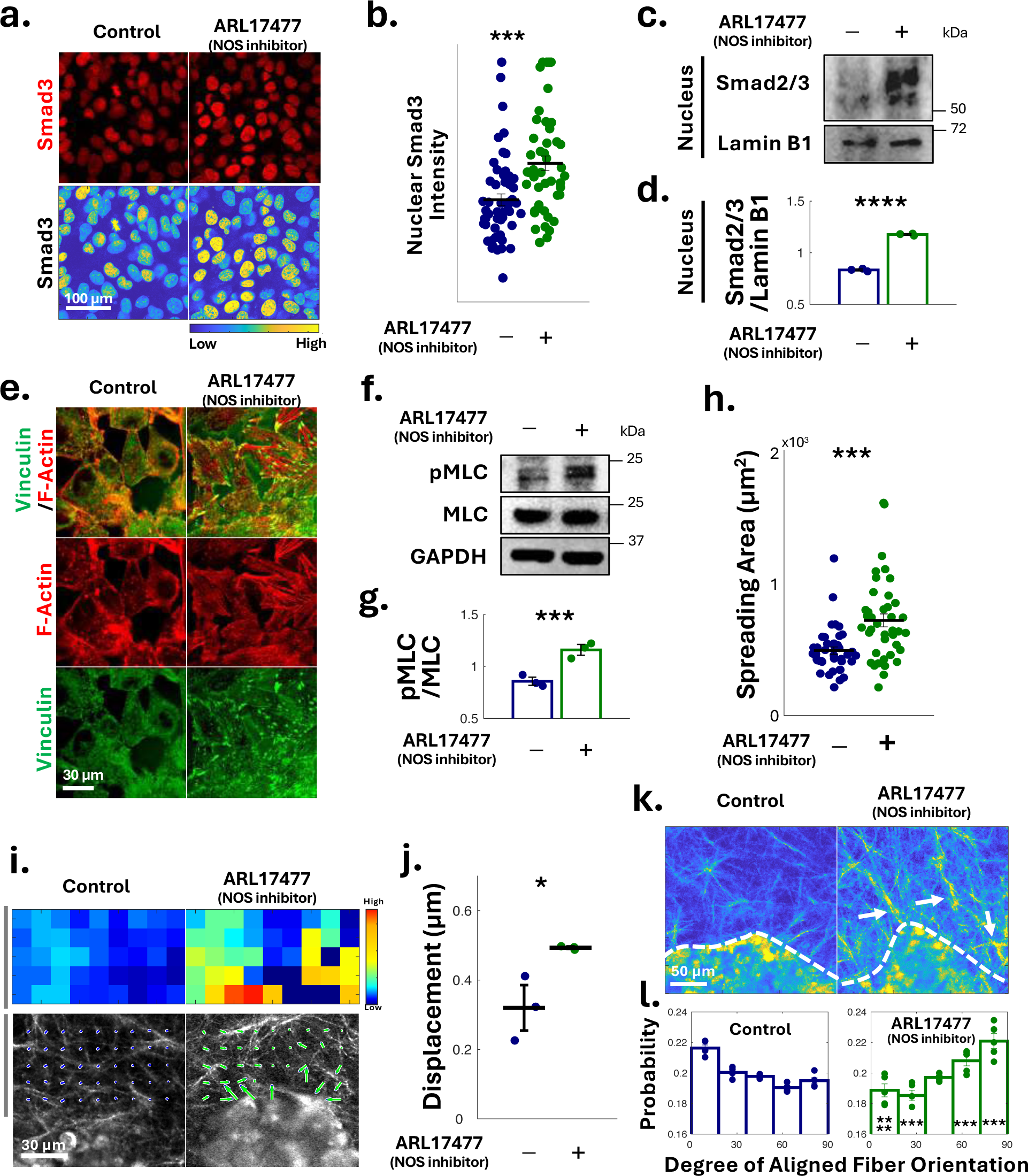
NO synthesis, promoted by PAX3-FOXO1, suppresses cell-ECM interaction. **(a)** Immunofluorescent and its pseudo-color images of Rh41 (FPRMS) cells stained for Smad3 with and without the treatment of a NOS inhibitor (ARL17477, 10 µM, 12 hours). **(b)** Intensity of nuclear Smad3 in Rh41 cells with and without the treatment of ARL17477. **(c)** Immunoblotting of nuclear fractionized proteins and **(d)** its quantification suggest that NOS inhibition increases nuclear localization of Smad3. Lamin B1 is a control of nuclear fractionized proteins. **(e)** Immunofluorescent images of Rh41 cell lines with and without the treatment of a NOS inhibitor (ARL17477, 10 µM, 12 hours), stained for F-actin (with phalloidin) and vinculin, showing focal adhesion. **(f)** Immunoblotting and **(g)** its quantification suggest that NOS inhibition increases phosphorylation of MLC, implying higher cell-ECM interaction. In **(d)** and **(g)**, error bars represent S.E. Statistical significance was assessed using two-sided Student’s t-tests: *P<0.05 (n = 3). **(h)** Cell spreading areas of Rh41 with and without NOS inhibition. **(i)** Collagen fiber displacement near spheroids with and without the treatment of ARL17477 (10 µM), monitored by CRM and analyzed by PIV. Top: pseudo-colored images showing the displacement degree of collagen fibers caused by spheroids. Bottom: Arrows indicate the direction and magnitude of collagen fiber displacement. **(j)** Quantification of collagen fiber displacement with and without ARL17477. In **(b), (h),** and **(j)**, each dot represents the value of a corresponding cell or spheroid. Lines are averages of each condition. Error bars represent S.E. Statistical significance was assessed using two-sided Student’s t-tests: *P<0.05 and ***P>0.005. **(k)** Collagen fiber alignment 16 hours post-gelation with Rh41 spheroids with and without the treatment of ARL17477. Dashed lines denote the spheroid edges. Arrows point to the fibers aligned perpendicularly to the spheroid edges. **(l)** Distribution of collagen fiber orientation around Rh41 spheroids with and without the NOS inhibitor. Dots represent individual spheroid values. Statistical differences of values between were assessed using two-sided Student’s t-test: ***P<0.005, and ****P <0.001.

### The PAX3-FOXO1 fusion gene determines anchorage-dependent tumor cell growth and metastatic potential

We hypothesized that the distinct cell-ECM interactions associated with the presence or absence of the PAX3-FOXO1 fusion gene would exhibit varying sensitivities to growth inhibition upon cell-ECM perturbation. FAK and Src are essential for forming focal adhesions, which link the cell cytoskeleton to the ECM^48–51^. These complexes facilitate bi-directional cell-ECM communication, which is crucial for regulating cell survival signals. Disruption of cell-ECM interactions by Src inhibition (Dasatinib) or FAK inhibition (Defactinib), respectively, resulted in diminished cell survival for FNRMS cells, which exhibit pronounced cell-ECM interaction, compared to FPRMS (**Figure 8a** and **Supplementary Figure 8a**). Similarly, inhibiting TGFβ signaling (LY2109761), an upstream regulator of cell-ECM interactions in our study, also resulted in growth inhibition for FNRMS but not FPRMS. Knockdown of PAX3-FOXO1 re-sensitized FPRMS cells to the growth-inhibiting effects of Src, FAK, and TGFβ inhibitors (**Figure 8b, Supplementary Figure 8b** and **c**). These results, consistent with higher TGFβ signaling-mediated cell-ECM interaction of FNRMS than FPRMS, suggest that targeted therapies to cell-ECM interaction would be effective to FNRMS but less to FPRMS. Its combination with the currently developing PAX3-FOXO1-targeted treatment, reducing its expression^52^, would successfully suppress FPRMS growth.

**Figure 8.**
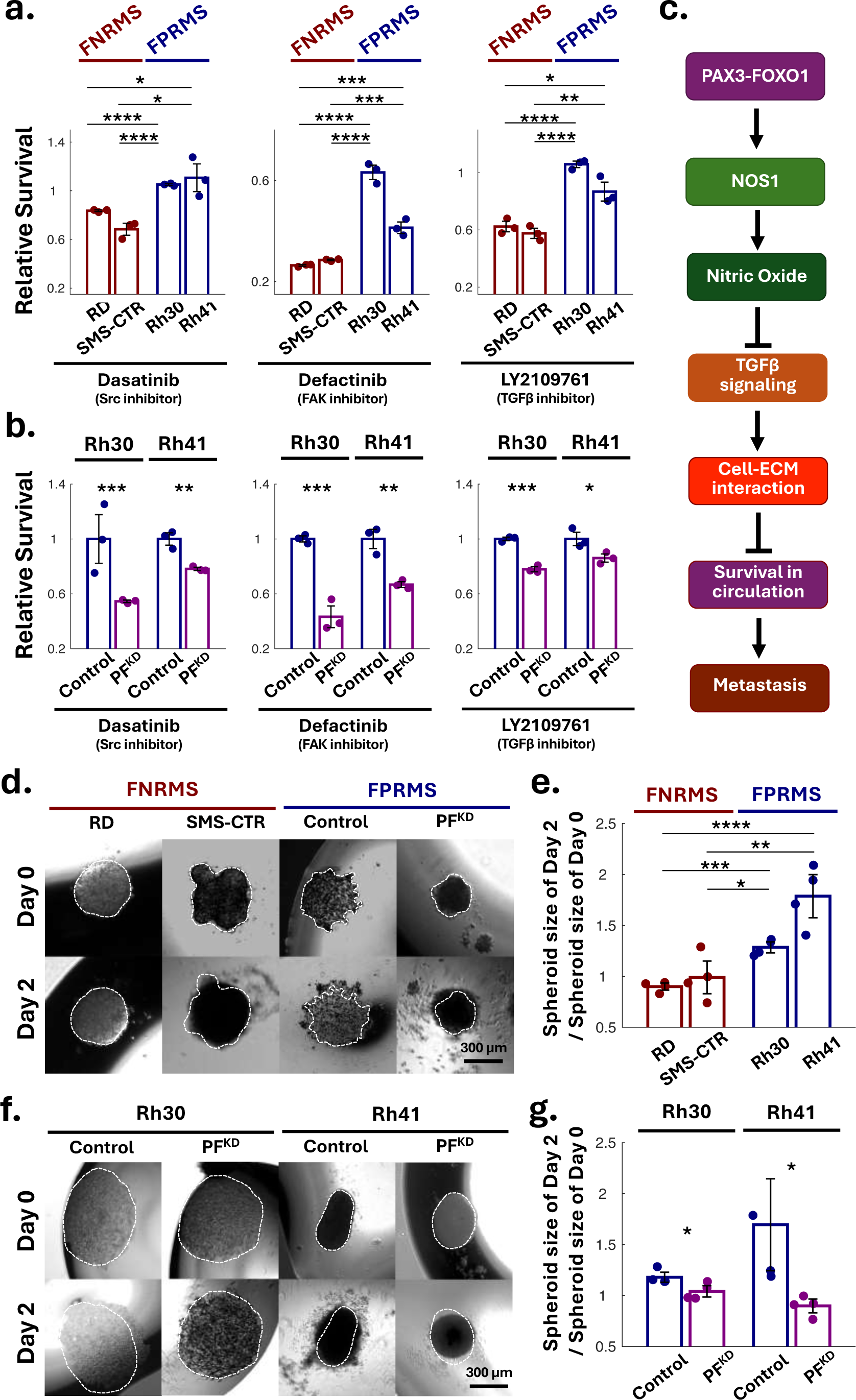
PAX3-FOXO1 modulates distinct cell-ECM interaction of FPRMS from FNRMS, determining their differential metastatic potentials. **(a)** PAX3-FOXO1 negatively regulates cell-ECM interaction and contributes to metastasis. **(b)** Higher resistance of FPRMS cell lines than FNRMS cell lines to pharmaceutical cell-ECM interaction perturbation and TGFβ inhibition. Relative survival of cells treated with Dasatinib (Src inhibitor, 10 nM, 1 day), Defactinib (FAK inhibitor, 10 µM, 1 day), and LY2109761 (TGFβ inhibitor, 1 mM, 1 day) compared to untreated cells. Cell survival was assessed by CCK8 assay. Error bars represent S.E. Statistical significance was assessed using a two-sided Student’s t-test: *P<0.05, **P<0.01, ***P<0.005, and ****P<0.001. **(c)** The PF knockdown decreases the resistance of cell-ECM interaction perturbation and TGFβ inhibition. **(d)** Suspended spheroids of FNRMS and FPRMS cells in media for 2 days via hang drop methods, no ECM interaction. **(e)** The growths of the suspended spheroids. Relative spheroid sizes (the ratios of spheroid sizes on day 2 to those on day 0) mean the degrees of their growth. **(f)** Suspended spheroids of control and PF^KD^ FPRMS cells for 2 days in the media via hang drop methods. **(g)** The growths of the suspended spheroids of PF^KD^ and control FPRMS cells. In **(d)** and **(f)**, dashed lines indicate the size of spheroid on day 0. In **(e)** and **(g)**, relative spheroid sizes (the ratios of spheroid sizes on day 2 to those on day 0) mean the degrees of their growth. Dots represent individual spheroid values. Asterisks (*) denote statistical differences, assessed by two-sided Student’s t-tests: *P<0.05, **P<0.01, ***P<0.005, and ****P<0.001.

FPRMS has been reported to have a higher metastatic potential compared to FNRMS ^1^. Cancer cells disseminate into blood or lymphatic vessels during metastasis, spreading through the circulation (**Figure 8c**). These circulating tumor cells (CTCs) may reprogram adherence dependency, suppressing cell-ECM interaction by deactivating YAP signaling and undergoing adherent-to-suspension transition (AST)^13^. CTCs can benefit from this reprograming, which enables them to resist to cell death within circulating system. We assessed the growth of the spheroids in the hanging drop condition without attaching to an ECM or solid surfaces, replicating the environment conditions that intravasated CTCs encounter. This setup effectively examines their ability to proliferate and grow independently of substrate attachment – a prerequisite for metastasis, potentially imparted by the PAX3-FOXO1 fusion gene. Indeed, the growth of FNRMS spheroids was inhibited in the hanging drop environment without ECM contact (**Figure 8d** and **e**). In contrast, FPRMS, less reliant on cell-ECM interaction, showed significant expansion over 48 hours and PF^KD^ nullified the expansion (**Figure 8f, g, Supplementary Figure 9a**, and **b**). Furthermore, PF^KD^ FRPMS cells exhibited higher nuclear localization of YAP, indicating YAP activation, and reduced expression of AST signature genes (**Supplementary Figure 9c** and **d**). These results indicate that the PAX3-FOXO1 status differentially regulates anchorage-dependent growth via distinct cell-matrix interactions, potentially contributing to the higher metastatic potential of FPRMS compared to FNRMS. The PAX3-FOXO1 fusion gene may confer metastatic competence by reprogramming anchorage dependence to sustain cancer cell growth and survival in ectopic environments, such as the circulatory system^13, 53^.

## Discussion

Our study investigated distinct characteristics between the two subtypes of RMS, FPRMS and FNRMS, to improve our understanding of the most common pediatric sarcoma and to develop better treatment strategies. Through sc-RNA seq data analysis, we confirmed that FPRMS and FNRMS signatures are enriched in different subgroups in the trajectory of normal human skeletal myogenesis, implying distinct differentiation arrests and signaling pathways of the RMS subtypes. We then discovered the differential expression of cell-ECM interaction-related genes between FPRMS- and FNRMS-like subgroups. Consistently, FNRMS exhibited higher expression of genes that regulates the interaction of cells with the ECM compared to FPRMS. Indeed, FNRMS cells demonstrated prominent molecular features and functional outcomes of cell-ECM interaction. Particularly, CRM enabled us to observe enhanced dynamic ECM reorganization in FNRMS compared to FPRMS, indicating increased cell-ECM interaction.

The oncogenic effect of the aberrant PAX3-FOXO1 transcription factor is thought to result from the transcriptional activation of multiple downstream target genes whose expression dysregulate signaling pathways that promote tumorigenesis. Several of these downstream effectors have activities that promote cancer development, such as stimulating proliferation, invasion, and inhibiting apoptosis. Some of these direct transcriptional targets of PAX3-FOXO1 include various RTKs, notably FGFR4^54^ and MET^55^. PAX3-FOXO1 also acts as a pioneer transcription factor that alters the cell epigenetic state to co-opt normal myogenic differentiation programs ^56^. Our findings expand on how the PAX3-FOX1 transcriptionally reprograms anchorage-dependency in RMS by downregulating genes involved in cell-matrix interactions. It does so by transcriptionally regulating a NO-mediated TGF-β signaling suppression that remodels that cytoskeletal architecture and disrupts focal adhesions to weaken cell-matrix association and potentiate cell survival.

The higher metastatic propensity of FPRMS can be attributed to the PAX3-FOXO1-mediated suppression of adhesion dependency. CTCs of FPRMS may benefit from this adhesion independence, allowing them to survive in the ECM-poor circulating system, in contrast to FNRMS cells that exhibit reduced survival in suspension. However, the initial step of metastasis, the invasion of cancer cells through adjacent stroma and the basement membrane, requires strong cell-ECM interaction^57^, which could be paradoxical to the higher metastatic potential of FPRMS that exhibits reduced interaction. Recently, Reginal *et al.* demonstrated that primary FPRMS cultures exhibit marked heterogeneity in PAX3-FOXO1 expression levels^58^. Additionally, they demonstrated that cells with lower expression levels showed higher adhesion to the ECM and invasiveness, which could initiate the early step of metastasis, while cells with higher expression may have a fitness advantage for surviving in ectopic environments during metastatic dissemination through the circulation. Thus, we need a better understanding of how FPRMS modulates the expression levels of PAX3-FOXO1 and the plasticity of cell-ECM interactions at the single-cell level to acquire different competencies during the complex, multi-step metastatic journey.

Our study also underscores the therapeutic implications of targeting cell-ECM interaction in RMS subtypes. The differential sensitivity of FNRMS and FPRMS to focal adhesion (e.g., FAK and Src) and TGFβ inhibitors suggests that disrupting cell-ECM interaction could be a viable strategy for FNRMS treatment. Additionally, combining cell-ECM interaction inhibitors with PAX3-FOXO1-targeted therapies may effectively suppress FPRMS growth. These proposed therapeutic strategies have clinical potential as targeted therapies, maximizing the curative effects and minimizing long-term sequelae potentially resulting from current conventional chemotherapies.

In conclusion, our findings highlight how gene fusion status impacts cell-ECM interaction, with FPRMS exhibiting reduced cell-ECM interaction compared to FNRMS. By elucidating the mechanisms by which PAX3-FOXO1 reprograms cell-ECM interaction, our study could lead to the development of targeted therapies that exploit their differential interaction, potentially improving clinical outcomes for RMS patients. Furthermore, the PAX3-FOXO1 reprogramming anchorage-dependence could shed light into the molecular mechanisms driving the higher metastatic propensity in FPRMS.

## Material

### Cell Culture

The FNRMS cell line, RD, was purchased from ATCC (CCL 136). The FNRMS cell line SMS-CTR and the FP-RMS cell lines, Rh30 and Rh41, were all obtained from the Childhood Oncology Group (COG). All cell lines were grown in high glucose DMEM supplemented with 20% Fetal Bovine Serum (FBS) and 1% penicillin/streptomycin. The PAX3-FOXO1 shRNA plasmid was kindly gifted from Dr. Mark E. Hatley (St. Jude). FPRMS cells Rh30 and Rh41 were transduced with lentiviruses expressing shRNA PAX3-FOXO1 (5’-CCGGTCTCACCTCAGAATTCAATTCCTCGAGGAATTGAATTCTGAGGTGAGATTTTTG-3’ and 5-’AATTCAAAAATCTCACCTCAGAATTCAATTCCTCGAGGAATTGAATTCTGAGGTGAGA-3’) and TET-pLKO-puro-scrambled used as a control. Both plasmids were co-transfected with packaging plasmids pMD2.G VSV-G envelope (#12259, Addgene) and psPAX2 packaging vector (#12260, Addgene) into 293T cells with Lipofectamine™ 3000 Transfection kit (Invitrogen # L3000001) according to manufacturer’s protocol. FPRMS cells were transduced with viral particles after filtering (0.45 µM). Selection of efficiently transduced cells was achieved by treatment with puromycin (2 μg/ml final concentration). To induce shRNAs, we incubated cells with doxycycline (50ng/ml) for 2 days.

### Single Cell Sequencing Analysis

Single-cell RNA sequencing (scRNA-seq) data was obtained from GSE147457, Wk6 to Yr41 and stored in the AnnData format, which is optimized for handling large-scale annotated data matrices. We used the `scanpy` library in Python for subsequent analysis. Initial quality control steps involved filtering out low-quality cells and genes. Cells with fewer than 500 genes were excluded to remove potential debris and doublets, respectively. To account for variations in sequencing depth between individual cells, the data was normalized by scaling the total counts for each cell to a fixed value. Following normalization, a log transformation was applied to stabilize the variance across genes, making the data more amenable to downstream analysis. Then, we identified highly variable genes, which are the most informative for distinguishing between different cell types.

We performed t-Distributed Stochastic Neighbor Embedding (t-SNE) to reduce the dimensionality of the dataset, capturing the most significant sources of variation. Cells were clustered using the Leiden algorithm with a resolution of 0.4, which detects communities in the data based on their nearest neighbors in the reduced dimensional space. Additionally, we performed Partition-based graph abstraction (PAGA) analysis to examine the connectivity between clusters. Student’s t-tests and false discovery rates were performed to identify differentially expressed genes in each cluster, or in other words, subgroup. Python algorithm workflow is attached as a Supplementary file.

### RNA Sequencing Analysis

Total RNA was extracted from PF^KD^ and control FPRMS cells using an RNeasy Mini Kit (Qiagen) following the manufacturer’s protocol. A Nanodrop ND-1000 spectrophotometer (NanoDrop Technologies) was used for DNA and mRNA quantification and qualification. RNA samples were quantified using a Qubit 2.0 Fluorometer (Life Technologies) and RNA integrity was checked using the Agilent TapeStation 4200 (Agilent Technologies). The RNA sequencing libraries were prepared using the NEBNext Ultra II RNA Library Prep Kit for Illumina using manufacturer’s instructions (New England Biolabs, Ipswich, MA, USA). Briefly, mRNAs were initially enriched with Oligo d(T) beads. Enriched mRNAs were fragmented for 15 minutes at 94°C. The first second strand cDNA were subsequently synthesized. cDNA fragments were end repaired and adenylated at 3’ ends, and universal adapters were ligated to cDNA fragments, followed by index addition and library enrichment by PCR with limited cycles. The sequencing libraries were validated on the Agilent TapeStation (Agilent Technologies, Palo Alto, CA, USA), and quantified by using a Qubit 2.0 Fluorometer (Life Technologies) as well as by quantitative PCR (KAPA Biosystems). The sequencing libraries were clustered on one flowcell lane. After clustering, the flowcell was loaded on the Illumina HiSeq instrument (4000 or equivalent) according to manufacturer’s instructions. The samples were sequenced using a 2×150bp Paired End configuration.

Raw sequence data (.bcl files) generated from the sequencer were converted into fastq files and de-multiplexed using Illumina’s bcl2fastq 2.17 software. One mismatch was allowed for index sequence identification. After investigating the quality of the raw data, sequence reads were trimmed to remove possible adapter sequences and nucleotides with poor quality using Trimmomatic v.0.36. The trimmed reads were mapped to the reference genome available on ENSEMBL using the STAR aligner v.2.5.2b. The STAR aligner uses a splice aligner that detects splice junctions and incorporates them to help align the entire read sequences. BAM files were generated as a result of this step. Unique gene hit counts were calculated by using the Counts feature from the Subread package v.1.5.2. Only unique reads that fell within exon regions were counted. After extraction of gene hit counts, the gene hit counts table was used for downstream differential expression analysis.

### Gene Set Enrichment and Gene Ontology Analysis (GSEA and GO analysis)

We identified differentially expressed genes (DEGs) in each subgroup from scRNA-seq results (GSE147457) and in each RMS subtype (GSE114621). Additionally, we normalized the expression levels of each gene in the FPRMS cell lines for both control and PF^KD^ conditions. We used the normalized values from Rh30 and Rh41 cell lines to calculate fold-change and adjusted p-values of each gene using Student’s t-test and the false discovery rate. Thresholds of fold change >0.5 or <-0.5 and adjusted p-value <0.05 were applied to identify DEGs. We used string version 12.0 (https://string-db.org/) for GO analysis and GSEA (https://www.gsea-msigdb.org/) to explore enriched signaling pathways in FNRMS-like clusters and PF^KD^ cells. For GSEA, we ranked DEGs based on fold changes.

### NeonGreen (NG)-Smad3 Live-cell sensor

NG-Smad3 Live-cell sensor was thankfully gifted from Dr. Lea Goentoro (Caltech). We transfected the plasmid using Lipofectamine™ 3000 Transfection kit (Invitrogen # L3000001) according to manufacturer’s protocol and selected the transfected cells using puromycin at a 2 µg/ml concentration. The transfected cells were replated on 6-well glass-bottomed plates and imaged on a Nikon Eclipse Ti2 inverted microscope equipped with a live cell chamber, maintaining humidity, body temperature, and 5% CO_2_. Images were acquired every 16 min for 1 hour, at least 10 minutes before TGFβ treatment and 50 minutes after the treatment. Using a Matlab-based algorithm, we assessed the Smad3 intensity in the nucleus over time and quantified TGFβ signaling activity.

### Immunoblotting

RMS cells were lysed from their respective drug-treated and knockdown cell lines and collected in RIPA buffer containing protease/phosphatase inhibitors (Halt Protease/Phosphatase Inhibitor Cocktail 100x). After centrifugation at 20,000g for 15 min to remove cellular debris, the supernatant was quantified using a BCA assay. Subsequently, the protein samples underwent SDS-PAGE using Mini-PROTEAN TGX Gels (4-20%, Bio-Rad) and were transferred onto polyvinylidene fluoride (PVDF) membranes. Membranes were then blocked in 3% BSA in TBST for 1 hour before being incubated overnight at 4°C with primary antibodies, including FOXO1, SMAD2/3, MLC2, pMLC2 (Ser 19), LAMINB1, and GAPDH, all diluted at 1:1000. Next, membranes were washed 3 times in TBST before exposure to HRP-conjugated IgG secondary antibodies (diluted at 1:2000-1:5000) for 1 hour at room temperature. Membranes were washed again 3 times with TBST for 5 min each to remove unbound secondary antibody. Protein bands were visualized using Clarity Western ECL substrate (Bio-Rad) using the iBright FL1500 imaging system.

### Immunofluorescent and DAF2-DA staining

RMS cells were seeded on collagen type I (10 µg/ml for 1 hour, Thermo #A1048301) coated coverslips. Following pertinent treatments with drug inhibitors, cells were fixed with 4% paraformaldehyde in PBS for 10 min, permeabilized with 0.1% Triton X-100 in PBS for 20 min and blocked with 1% BSA in PBS for 45 min. Coverslips were incubated for 2 hours at room temperature with primary antibodies diluted in blocking solution (1:100-1:250 dilution range) and then washed 3 times with PBS to remove unbound antibodies. Coverslips were then incubated with Alexa-488 or Alexa-594 secondary antibodies (Invitrogen) at a concentration of 5 µg/mL in blocking solution for 1 hour at room temperature. The coverslips were then washed 3 times with PBS to remove unbound secondary antibodies and mounted on cover glass after counterstaining with DAPI (Prolong Gold antifade reagent; Invitrogen). Epifluorescence and confocal imaging were performed on a Nikon Eclipse Ti2 inverted microscope. We analyzed the images using a customized MATLAB script to read out the intensity score of each pixel in regions of interest, indicating the expression levels of corresponding proteins. Statistical analysis was performed using two-sided Student’s t-tests.

### Cell survival assay (CCK8 assay)

Cell viability was quantitatively assessed with a colorimetric assay using the Cell Counting Kit-8 (CCK-8, Sigma Aldrich #96992). In brief, 100 μL of cell suspension (5000 cells/well) was dispensed in a 96-well plate. The plate was pre-incubated for 24 hours in a humidified incubator (37C, 5% CO_2_) followed by the addition of drug inhibitor treatments into the media at the appropriate concentrations. The plate was further incubated for another 48 hours in the incubator and subsequently cell viability was assessed by adding 10 μL of CCK-8 solution into each well. The plate was incubated for 4 hours in the incubator and then the absorbance was measured at 450 nm using a microplate reader. Control wells with media only were used for normalization by subtracting the background signal of media-only wells from each absorbance measurement in cell-containing wells. Triplicates were used for each control and drug-treated condition, e.g. Dasatinib (Src inhibitor), Defactinib (FAK inhibitor), and LY2108761 (TGFβ inhibitor). Relative survival is reported as the average normalized absorbance measurement.

### Traction Force Microscopy

Traction force microscopy (TFM) experiment was performed on 0.6 kPa and 12.7 kPa high refractive index silicone gel by mixing two parts of Q-gel 920 A and B (Q-gel, CHT) in ratios 1:1 and 1:2 respectively as previously described ^59, 60^. The silicone Q gel was spin-coated on a 35 mm glass-bottom dish (#1 cover glass, from MatTek corporation), and cured at 80°C for 2 hours, and stored in phosphate buffer saline (PBS). To visualize gel deformation, the TFM gel surface was functionalized with (3-aminopropyl) triethoxysilane (APTES, Sigma-Aldrich, #440140) and covalently bonded with 40 nm diameter carboxylated far-red fluorescent beads (excitation/emission 690/720 nm, from Invitrogen) at a density of 1 bead /µm2 using 1-Ethyl-3-(3-dimethyl aminopropyl) carbodiimide (EDC, Sigma-Aldrich, #39391). As an adhesion protein, Collagen-II (Southern Biotech, # 35209) was coated in two different quantities, 0.5 µg/ml for sparse and 100 µg/ml for dense collagen density.

Cells were plated on the sparsely and densely collagen-II-coated gel substrates. After seeding for 4 hours, single-shot brightfield images of live cells and beads images at 642 nm were taken using a 60x objective total internal reflection fluorescence (TIRF) microscope (optoTIRF, CAIRN Research, Faversham ME13 8UP, UK) housed in a Nikon Ti-S microscope. The relaxed bead images as reference images for TFM were obtained after removing the cells using 0.5 ml of 10% bleach.

Using the bead and cell images, traction force reconstruction was performed as previously described^60, 61^. To calculate the displacement field, an image cross-correlation-based particle tracking velocimetry method (cPTVR) was used by comparing bead images before and after the cell is present on the substrate. The force reconstruction was done using the Fast Boundary Element Method (FastBEM) with L2-norm-based regularization. The regularization parameter was chosen based on the L-curve, L-corner method.

### Spheroid

RMS single-cell suspension was used for hanging drop cultures by detaching adherent cancer cell cultures at ∼70% confluence, followed by exposure to a 0.05% trypsin-EDTA solution. A dilution was then performed to allow for the seeding of 20,000 cells per 20 µl drop of cell culture media, which were deposited across the lid of a 60mm petri dish. The lid was carefully inverted and placed over the PBS-filled bottom chamber of the petri dish. The hanging drops were maintained in a 37°C, 5% CO_2_ incubator for 48 hours until compact aggregated spheroids formed. For the embedding of spheroids into a collagen-based ECM matrix, a 3 mg/mL stock of rat tail collagen I solution (Gibco) was used to prepare a total volume of 1 mL of pH-buffered collagen gel solution at a final density of 2 mg/mL. In a sterile tube, 100 μL of 1M HEPES, 100 μL of 10x DMEM, and 17 μL of 1N NaOH were mixed with 800 μL of collagen I solution. The collagen I solution was kept on ice to prevent polymerization. The spheroids formed in hanging drops were then collected in growth media, centrifuged at low speed (100g for 2 min), resuspended in growth media, and embedded in the collagen I solution at a 1:3 ratio (cells : gel mixture). LY2109761 (10 µM) and ARL17477 (10 µM) were also added to the mixture when investigating theeffect of these drugs. The resulting cell-encapsulating three-dimensional (3D) gel mixture (200 µl) was deposited in wells of a glass-bottom 24-well plate. The plate was incubated for 1 hour to allow the 3D gel : cell mixture to polymerize, and media was then added on top.

### Confocal reflection microscopy (CRM)

Live cell imaging with confocal reflection microscopy (CRM) for 16 hours was conducted to visualize collagen re-organization and remodeling of the collagen fiber network by RMS spheroids as a proxy for cell-ECM interactions. CRM imaging relies on the reflection (back-scattering) of light from structures within a biological sample that have differences in refractive indices. Collagen, being a fibrous protein with a higher refractive index than surrounding medium or cells, can produce a strong reflectance signal in confocal microscopy. This reflectance signal can be used to visualize dynamic collagen re-organization, morphology, and remodeling as the RMS tumor spheroids interact with the 3D matrix. In short, CRM was used to capture images every 3 hours. Samples of RMS tumor spheroids within the 3D matrix were imaged using an inverted Nikon Eclipse Ti2 microscope with a 20x objective lens. A continuous wave laser at 488 nm was employed for imaging the spheroids. Despite collecting the 475–495 nm bandwidth centered on the 488-reflection using 488 laser sources for CRM, the guiding mirror (45° reflection mirror) was eliminated to enable light reflection on the main beam splitter (MBS 80/20) at a 90° angle, maximizing excitation light of the same wavelength that illuminated the samples. The pinhole size for image acquisition was set at 1 AU (5 µm). To improve signal-to-noise and image quality, a dwelling time of 22.3 μs was used.

### Collagen Alignment Analysis

We utilized Particle Image Velocimetry (PIV) to analyze collagen displacement for 3 hours within the CRM images. Using the `matpiv` function in MATLAB, we extracted two time-series images. The interrogation window size was set to 25 µm, and the analysis was performed with a 50% overlap ratio to ensure dense vector fields. A single-pass correlation method was employed to compute the displacement vectors. The outputs included the grid coordinates `x` and `y`, and the corresponding velocity components `u` and `v`, corresponding to the magnitude of the displacement in x and y directions.

Furthermore, we developed a MATLAB-based algorithm to analyze the directions of collagen fibers in CRM images. First, we applied a Gaussian filter to smooth the image and reduce noise. For collagen segmentation, we utilized the Sobel filter in both horizontal and vertical directions. Horizontal edge detection was achieved by directly applying the Sobel filter, while vertical edge detection was obtained by using the transposed Sobel filter. To isolate the fibers in the images, a threshold was determined based on the mean and standard deviation of the image intensities, and a binary mask was created to highlight regions exceeding this threshold. The masked image was then used to calculate the angles of collagen alignment with respect to the x-axis, corresponding to the edges of nearby spheroids.

---

**Supplementary Figure 1.**
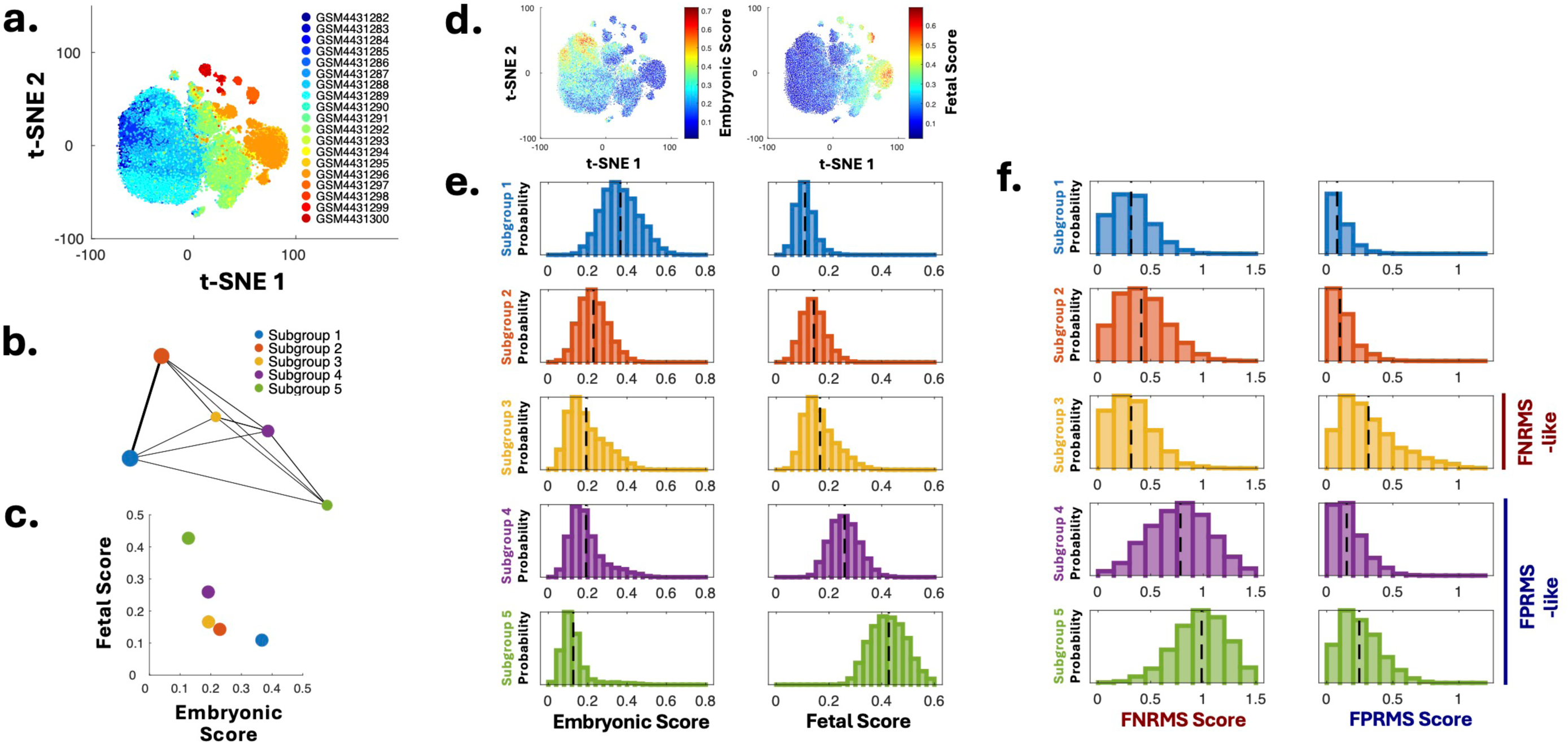
**(a)** t-SNE rendering single-cell RNA (scRNA) sequencing of skeletal muscle tissues obtained from human prenatal development and juveniles/adults from GSE147457, indicating cells from different samples. **(b)** Partition-based graph abstraction (PAGA), reconciling clustering with myogenic differentiation trajectory inference. **(c)** Fetal score, the expression levels of highly expressed genes in Subgroup 5, highly enriched in fetal tissues, and embryonic score, those of highly expressed genes in Subgroup 1, highly enriched embryonic tissues, of each subgroup. Higher fetal scores and lower embryonic scores indicate a later stage in differentiation. **(d)** Decreasing embryonic and increasing embryonic scores from Subgroups1 to 5. **(e)** Histogram of embryonic (left) and fetal (right) scores in each subgroup. (**f)** Histogram of FNRMS (left) and FPRMS (right) scores in each subgroup. In **(e)** and **(f)**, dashed lines indicate averages of each score of each subgroup.

**Supplementary Figure 2.**
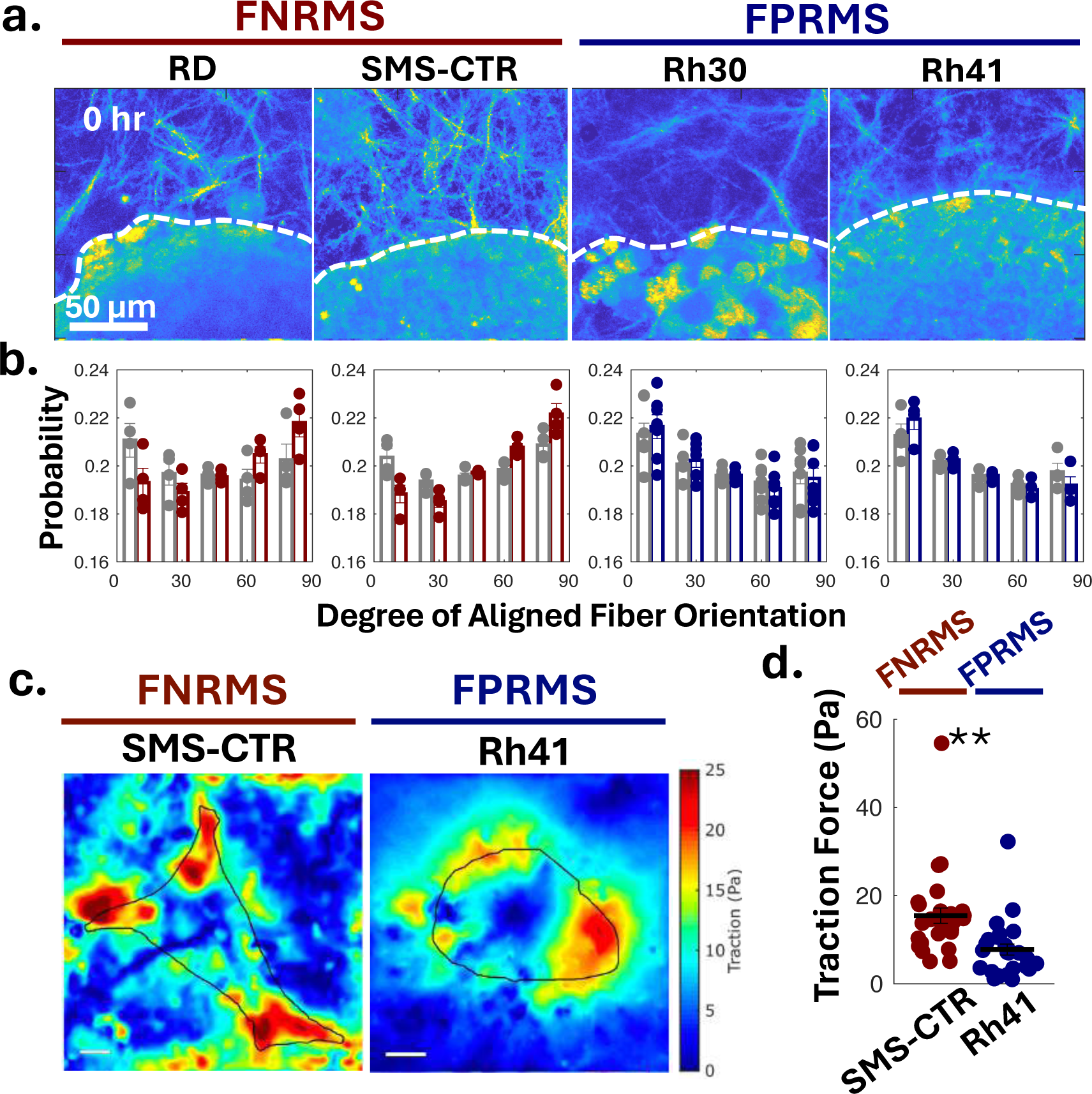
**(a)** Collagen fiber alignment 0 hours post-gelation with spheroids. Dashed lines denote spheroid edges. **(b)** Distribution of collagen fiber orientation around RMS cell spheroids at 0 hour (gray) and 16 hours (color, Fig. 2f). Angles at 90° represent perpendicular alignment to spheroid edges; 0° represents parallel alignment. Dots represent individual spheroid values. **(c)** Traction force microscopic images of SMS-CTR (FNRMS) and Rh41 (FPRMS) cells. **(d)** Traction force of SMS-CTR (FNRMS) and Rh41 (FPRMS) cells. Each dot indicates the value of each cell. Lines are averages of each group. Error bars are S.E. Statistical significance was assessed using two-sided Student’s t-tests: **P<0.01.

**Supplementary Figure 3.**
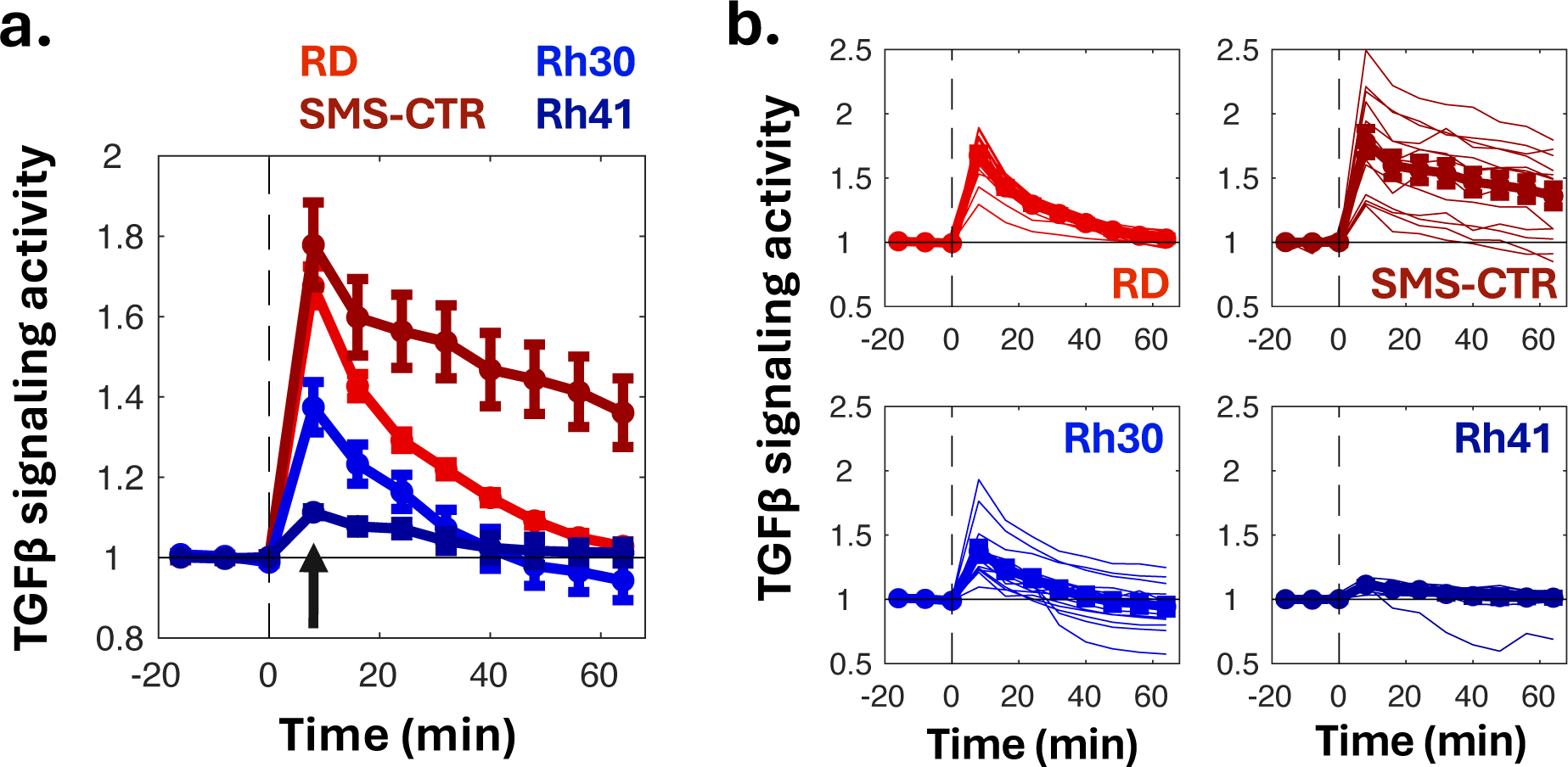
Time-course TGFβ signaling activity of RMS cell lines, assessed by dynamic Smad3 nuclear localization. **(a)** Averages of each cell line and **(b)** faint lines, indicating each cell in the cell line (right) Dashed lines indicate the time point of TGFβ treatment. An arrow indicates the value of Fig. 3c.

**Supplementary Figure 4.**
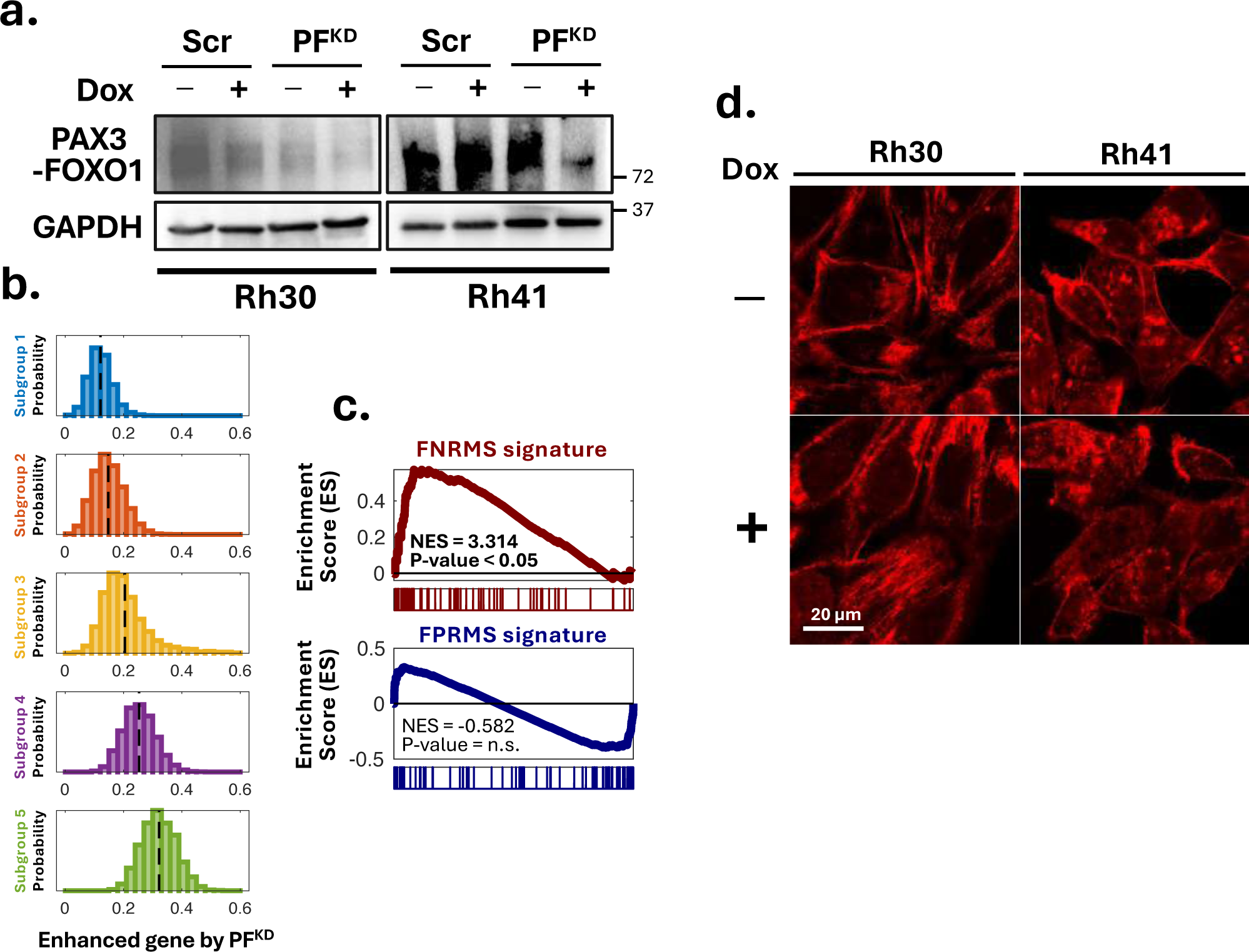
**(a)** PAX3-FOXO1 expression levels of scramble and PF^KD^ cells with and without doxycycline (Dox) induction. **(b)** Histogram of the average expression levels of genes enhanced by PF^KD^ in each cell of each subgroup identified by scRNA seq of developing skeletal muscle tissues. Dashed lines indicate averages of each subgroup. **(c)** GSEA of genes enhanced by PF^KD^ with FNRMS (top) and FPRMS (bottom) signatures as the reference gene sets. The genes enhanced by PF^KD^ exhibit statistically significant overlap with FNRMS signatures, not FPRMS signatures (P-value = no significance). **(d)** F-actin stained by phalloidin in scramble FPRMS cells displayed cortical actin, indicating reduced cell-ECM interaction.

**Supplementary Figure 5.**
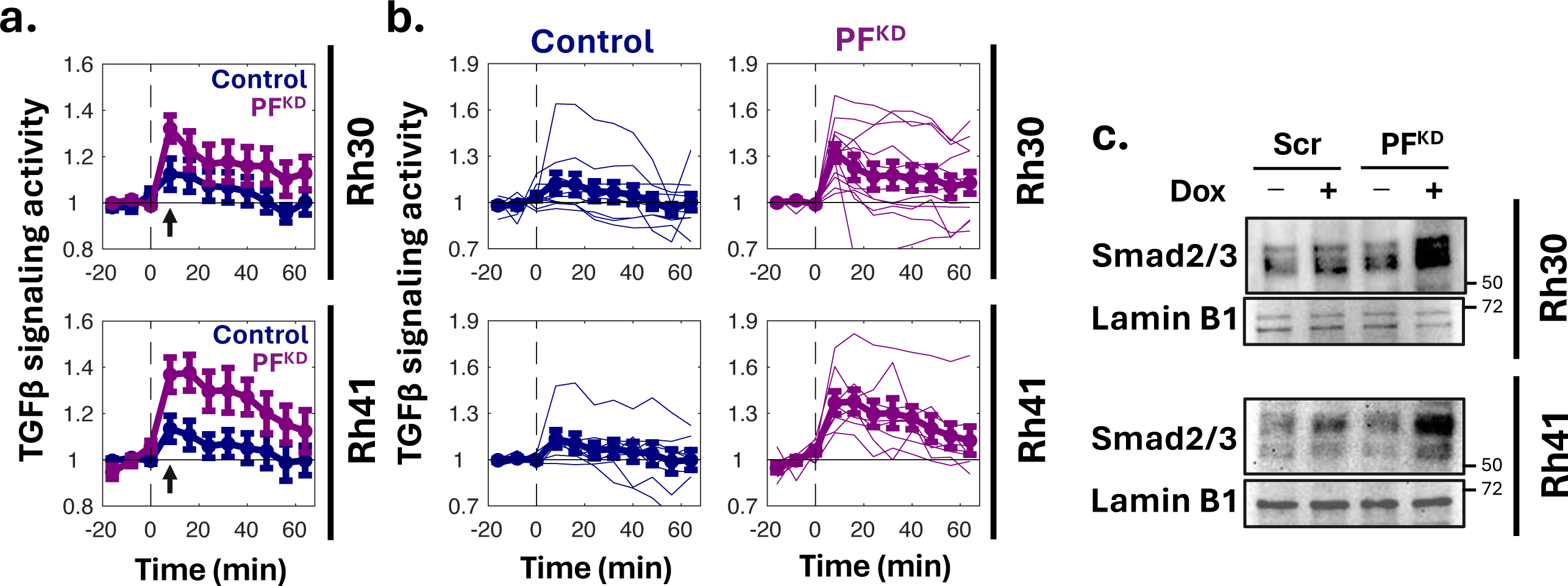
**(a)** Averaged time-course TGFβ signaling activity of PF^KD^ (purple) and control (blue) FPRMS cells, assessed by dynamic Smad3 nuclear localization in each cell line. **(b)** Faint lines indicate each cell in the cell line. Dashed lines indicate the time point of TGFβ treatment. Arrows indicate the values of Fig. 4i. **(c)** Higher Smad3 nuclear localization of PF^KD^ FPRMS cells compared to control scramble cells.

**Supplementary Figure 6.**
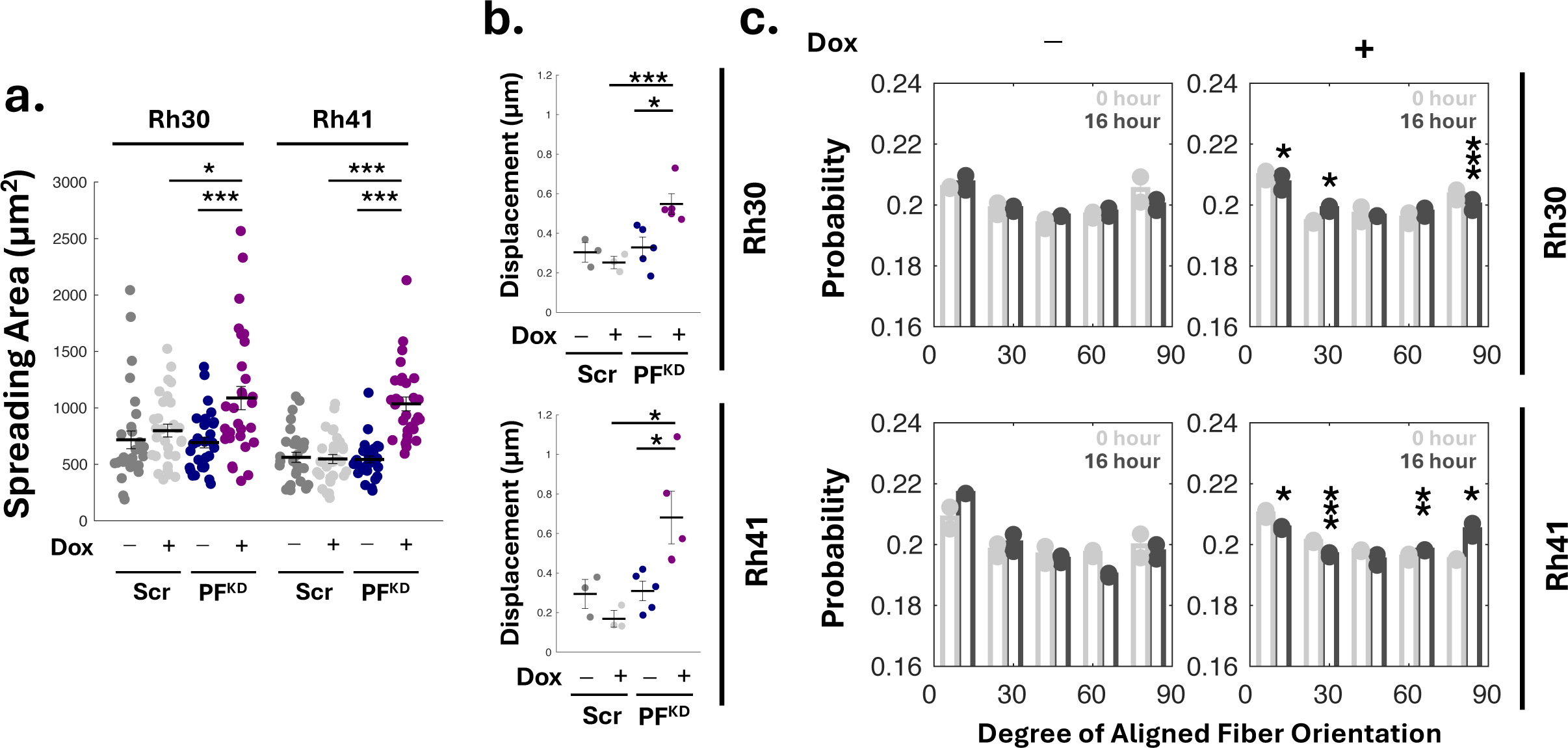
**(a)** Pronounced cell spreading areas of PF^KD^ cells compared to control scramble cells. **(b)** Displacement of collagen fibers around spheroids of scrambled and PF^KD^ FRPMS cells, Rh30 (top) and Rh41 (bottom), assessed by CRM. In **(a)** and **(b)**, each dot denotes the value of a corresponding cell and lines are the averages of each cell line. Error bars are S.E. Statistical significance was assessed using two-sided Student’s t-tests: *P<0.05 and ***P<0.005. **(c)** Distribution of collagen fiber orientation around the spheroids of scramble FPRMS cells at 0 and 16 hours. At 16 hours, statistical significances between Dox-induced scramble and PF^KD^ cells (**Figure 5e**) were assessed by two-sided Student’s t-tests: P*<0.05, P**<0.01, and P***<0.005.

**Supplementary Figure 7.**
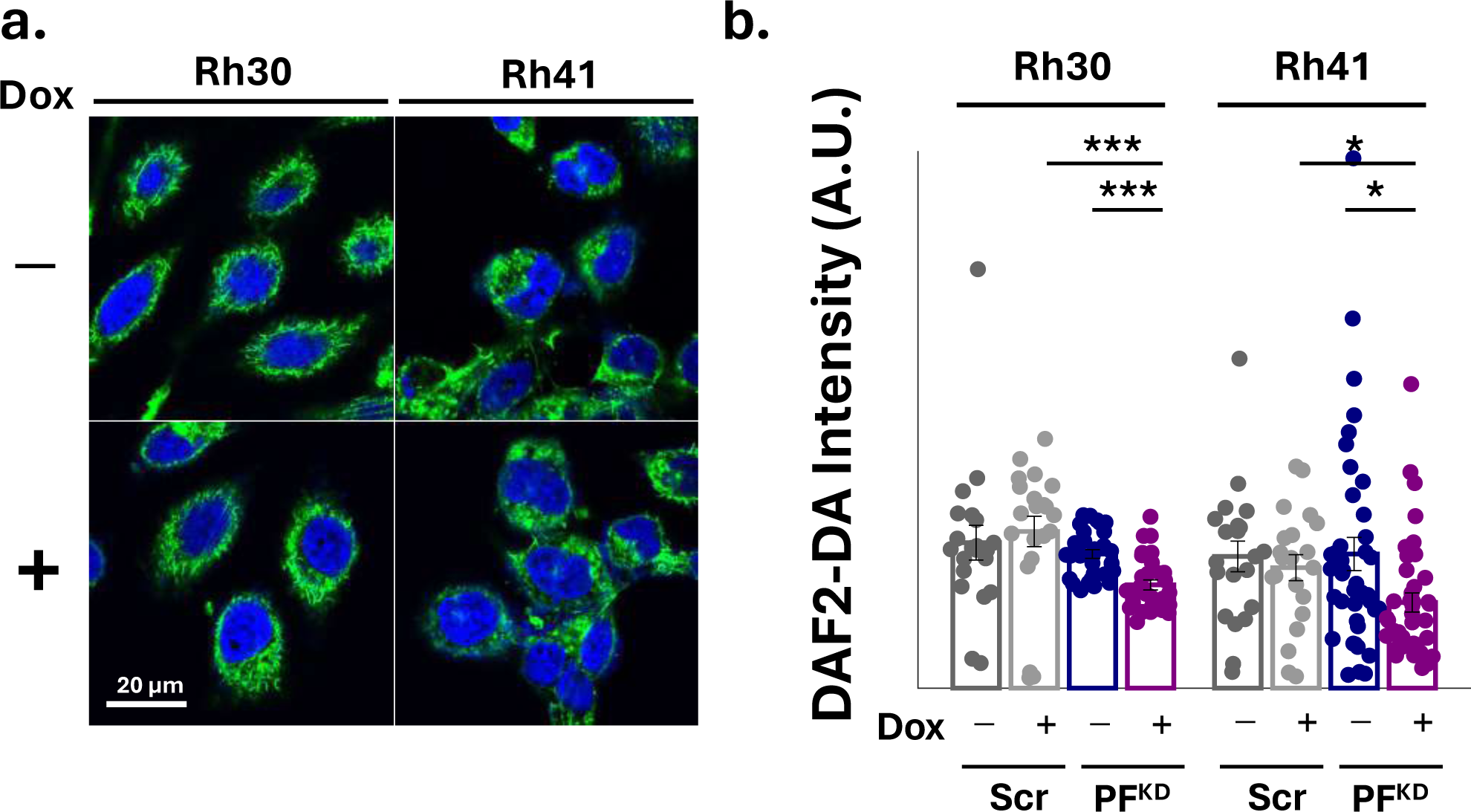
**(a)** DAF2-DA staining of scramble FPRMS cells, indicating their intracellular NO concentration. **(b)** Quantification of DAF2-DA, corresponding to NO concentration, of scramble and PF^KD^ FPRMS (**Figure 6e** and **f**). Dots represent individual the values of corresponding cells. Statistical differences were assessed by two-sided Student’s t-tests: *P<0.05, and ***P<0.005.

**Supplementary Figure 8.**
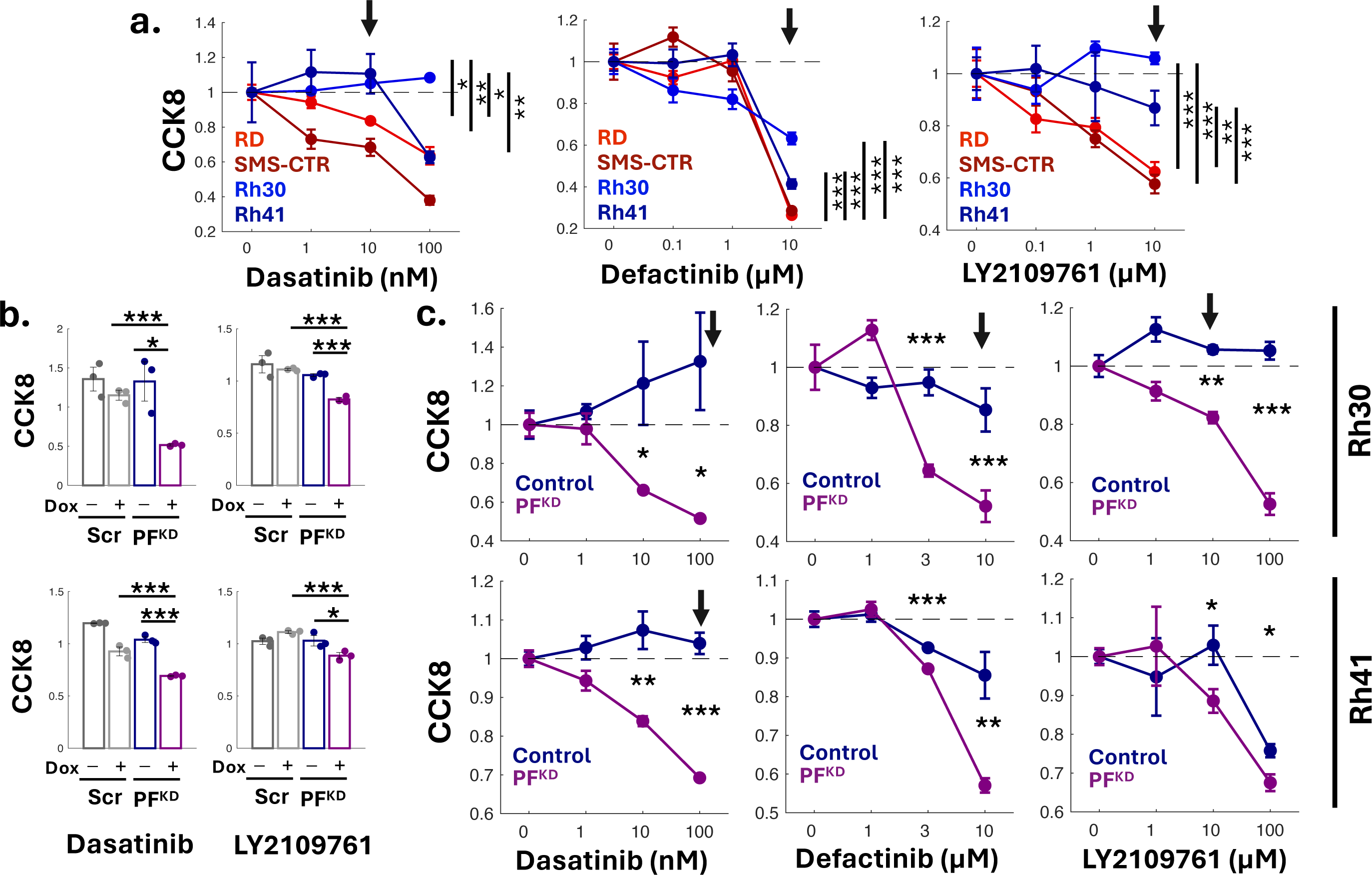
**(a)** Higher resistance of FPRMS than FNRMS to pharmaceutical cell-ECM perturbation (Src inhibitor, Dasatinib; FAK inhibitor, Defactinib) and TGFβ inhibition (TGFβ inhibitor, LY219761). Statistical significance of cell survivals in the arrow marked drug concentration was assessed by two-sided Student’s t-tests: P*<0.05, P**<0.01, P***<0.005, and P****<0.001. **(b)** Cell survival of scramble and PF^KD^ FPRMS cells, Rh30 (top) and Rh41 (bottom), upon the treatment of Dasatinib (10 µM) and LY2109761 (10 µM). Statistical significance was assessed by two-sided Student’s t-tests: P*<0.05, and P***<0.005. **(c)** Higher sensitivity of PF^KD^ FPRMS cells than control cells to pharmaceutical cell-ECM perturbation (Src inhibitor, Dasatinib; FAK inhibitor, Defactinib) and TGFβ inhibition (TGFβ inhibitor, LY219761). Statistical significance of cell survival was assessed by two-sided Student’s t-tests: P*<0.05, P**<0.01, P***<0.005, and P****<0.001. Arrows indicate the concentration used in main figures.

**Supplementary Figure 9.**
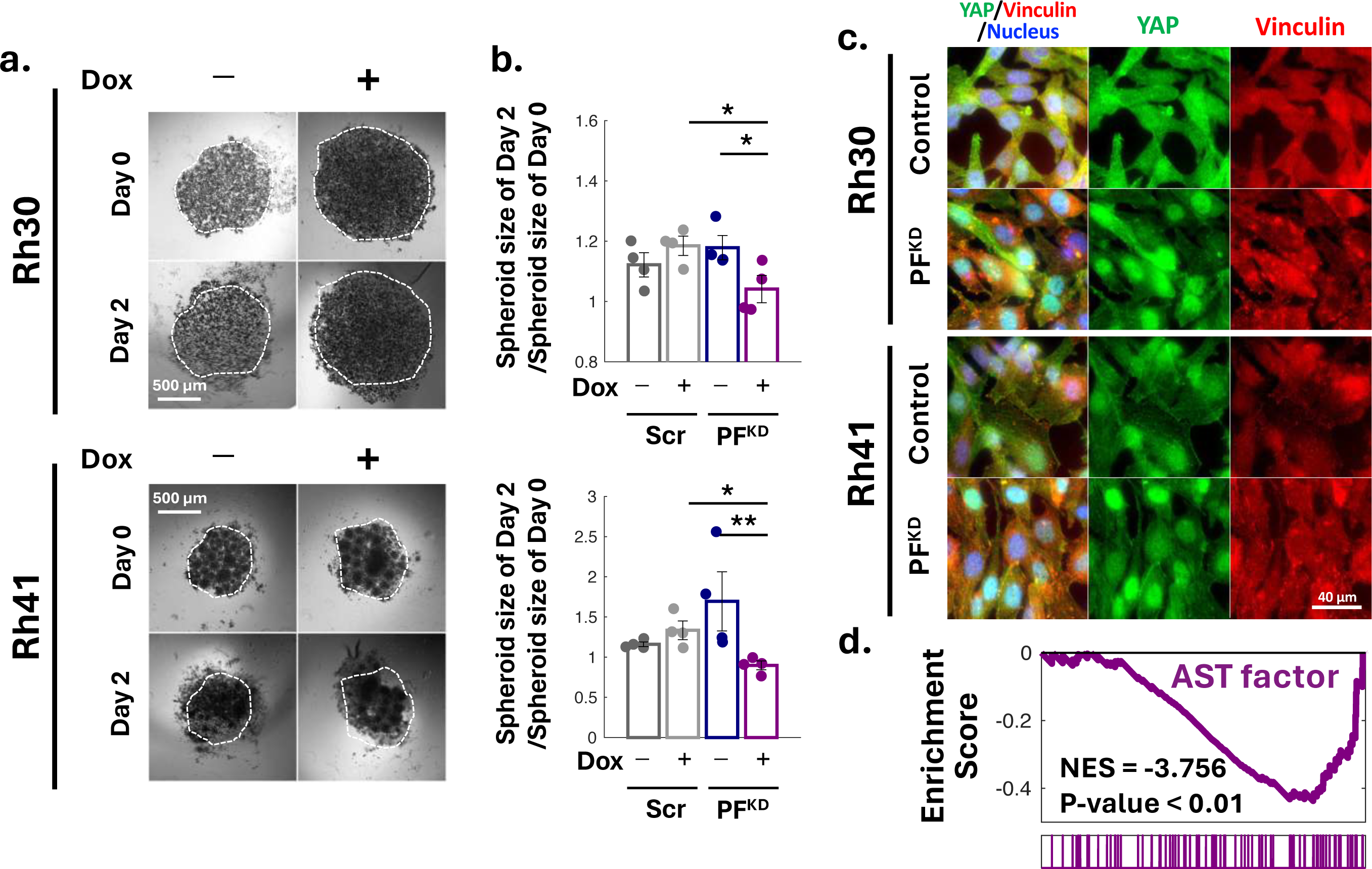
**(a)** Suspended spheroids of scramble FPRMS cells in the media for 2 days via hang drop methods, no ECM interaction. Dashed lines indicate the size of spheroid on day 0. **(b)** Growth of the suspended spheroids of scramble and PF^KD^ FPRMS (**Figure 8f** and **g**). Relative spheroid sizes (the ratios of spheroid sizes on day 2 to those on day 0) mean the degrees of their growth. Relative spheroid sizes (the ratios of spheroid sizes on day 2 to those on day 0) mean the degrees of their growth. Dots represent individual spheroid values. Statistical difference was assessed by two-sided Student’s t-tests: *P<0.05, and **P<0.01. **(c)** Immunofluorescent images of PF^KD^ and control cells FPRMS cells stained for YAP and vinculin, Higher nuclear localization of YAP in PF^KD^ cells means their higher YAP activation than control cells. **(d)** GSEA suggests that reduced expression of AST markers in PF^KD^ FPRMS cells compared to control cells.

## Notes

### Competing Interest Statement

The authors have declared no competing interest.

## References

1. Skapek, S.X. et al. Rhabdomyosarcoma. Nat Rev Dis Primers 5, 1 (2019).

2. Zarrabi, A. et al. Rhabdomyosarcoma: Current Therapy, Challenges, and Future Approaches to Treatment Strategies. Cancers (Basel) 15 (2023).

3. Chen, C., Dorado Garcia, H., Scheer, M. & Henssen, A.G. Current and Future Treatment Strategies for Rhabdomyosarcoma. Front Oncol 9, 1458 (2019).

4. Wei, Y. et al. Single-cell analysis and functional characterization uncover the stem cell hierarchies and developmental origins of rhabdomyosarcoma. Nat Cancer 3, 961–975 (2022).

5. Pickup, M.W., Mouw, J.K. & Weaver, V.M. The extracellular matrix modulates the hallmarks of cancer. EMBO Rep 15, 1243–1253 (2014).

6. Winkler, J., Abisoye-Ogunniyan, A., Metcalf, K.J. & Werb, Z. Concepts of extracellular matrix remodelling in tumour progression and metastasis. Nat Commun 11, 5120 (2020).

7. Cox, T.R. The matrix in cancer. Nat Rev Cancer 21, 217–238 (2021).

8. Osses, N. & Brandan, E. ECM is required for skeletal muscle differentiation independently of muscle regulatory factor expression. Am J Physiol Cell Physiol 282, C383–394 (2002).

9. Choi, Y.J. et al. Muscle-derived extracellular matrix on sinusoidal wavy surfaces synergistically promotes myogenic differentiation and maturation. J Mater Chem B 6, 5530–5539 (2018).

10. Melo, F., Carey, D.J. & Brandan, E. Extracellular matrix is required for skeletal muscle differentiation but not myogenin expression. J Cell Biochem 62, 227–239 (1996).

11. Rozario, T. & DeSimone, D.W. The extracellular matrix in development and morphogenesis: a dynamic view. Dev Biol 341, 126–140 (2010).

12. Thorsteinsdottir, S., Deries, M., Cachaco, A.S. & Bajanca, F. The extracellular matrix dimension of skeletal muscle development. Dev Biol 354, 191–207 (2011).

13. Huh, H.D. et al. Reprogramming anchorage dependency by adherent-to-suspension transition promotes metastatic dissemination. Mol Cancer 22, 63 (2023).

14. Xi, H. et al. A Human Skeletal Muscle Atlas Identifies the Trajectories of Stem and Progenitor Cells across Development and from Human Pluripotent Stem Cells. Cell Stem Cell 27, 158–176 e110 (2020).

15. Traag, V.A., Waltman, L. & van Eck, N.J. From Louvain to Leiden: guaranteeing well-connected communities. Sci Rep 9, 5233 (2019).

16. Gryder, B.E. et al. PAX3-FOXO1 Establishes Myogenic Super Enhancers and Confers BET Bromodomain Vulnerability. Cancer Discov 7, 884–899 (2017).

17. Drummond, C.J. et al. Hedgehog Pathway Drives Fusion-Negative Rhabdomyosarcoma Initiated From Non-myogenic Endothelial Progenitors. Cancer Cell 33, 108–124 e105 (2018).

18. Searcy, M.B. et al. PAX3-FOXO1 dictates myogenic reprogramming and rhabdomyosarcoma identity in endothelial progenitors. Nat Commun 14, 7291 (2023).

19. Berrier, A.L. & Yamada, K.M. Cell-matrix adhesion. J Cell Physiol 213, 565–573 (2007).

20. Wozniak, M.A., Modzelewska, K., Kwong, L. & Keely, P.J. Focal adhesion regulation of cell behavior. Biochim Biophys Acta 1692, 103–119 (2004).

21. Zhao, J. & Guan, J.L. Signal transduction by focal adhesion kinase in cancer. Cancer Metastasis Rev 28, 35-49 (2009).

22. Artym, V.V. & Matsumoto, K. Imaging cells in three-dimensional collagen matrix. Curr Protoc Cell Biol Chapter 10, Unit 10 18 11-20 (2010).

23. Brightman, A.O. et al. Time-lapse confocal reflection microscopy of collagen fibrillogenesis and extracellular matrix assembly in vitro. Biopolymers 54, 222–234 (2000).

24. Foty, R. A simple hanging drop cell culture protocol for generation of 3D spheroids. J Vis Exp (2011).

25. Petitjean, L. et al. Velocity fields in a collectively migrating epithelium. Biophys J 98, 1790–1800 (2010).

26. Raffel, M. et al. Particle image velocimetry: a practical guide. (springer, 2018).

27. Provenzano, P.P. et al. Collagen reorganization at the tumor-stromal interface facilitates local invasion. BMC Med 4, 38 (2006).

29. Mittal, N. & Han, S.J. High-Resolution, Highly-Integrated Traction Force Microscopy Software. Curr Protoc 1, e233 (2021).

29. Liu, A.P., Chaudhuri, O. & Parekh, S.H. New advances in probing cell-extracellular matrix interactions. Integr Biol (Camb) 9, 383–405 (2017).

30. Style, R.W. et al. Traction force microscopy in physics and biology. Soft Matter 10, 4047–4055 (2014).

31. Verrecchia, F. & Mauviel, A. Transforming growth factor-beta signaling through the Smad pathway: role in extracellular matrix gene expression and regulation. J Invest Dermatol 118, 211–215 (2002).

32. Ignotz, R.A. & Massague, J. Transforming growth factor-beta stimulates the expression of fibronectin and collagen and their incorporation into the extracellular matrix. J Biol Chem 261, 4337–4345 (1986).

33. Richardson, L., Wilcockson, S.G., Guglielmi, L. & Hill, C.S. Context-dependent TGFbeta family signalling in cell fate regulation. Nat Rev Mol Cell Biol 24, 876–894 (2023).

34. Massague, J. TGFbeta signalling in context. Nat Rev Mol Cell Biol 13, 616–630 (2012).

35. Schmitt-Ney, M. & Camussi, G. The PAX3-FOXO1 fusion protein present in rhabdomyosarcoma interferes with normal FOXO activity and the TGF-beta pathway. PLoS One 10, e0121474 (2015).

36. Frick, C.L., Yarka, C., Nunns, H. & Goentoro, L. Sensing relative signal in the Tgf-beta/Smad pathway. Proc Natl Acad Sci U S A 114, E2975–E2982 (2017).

37. Zieba, A. et al. Intercellular variation in signaling through the TGF-beta pathway and its relation to cell density and cell cycle phase. Mol Cell Proteomics 11, M111 013482 (2012).

38. Hanna, J.A. et al. PAX3-FOXO1 drives miR-486-5p and represses miR-221 contributing to pathogenesis of alveolar rhabdomyosarcoma. Oncogene 37, 1991–2007 (2018).

39. Kassianidou, E., Hughes, J.H. & Kumar, S. Activation of ROCK and MLCK tunes regional stress fiber formation and mechanics via preferential myosin light chain phosphorylation. Mol Biol Cell 28, 3832–3843 (2017).

40. Totsukawa, G. et al. Distinct roles of MLCK and ROCK in the regulation of membrane protrusions and focal adhesion dynamics during cell migration of fibroblasts. J Cell Biol 164, 427–439 (2004).

41. Goligorsky, M.S. et al. Nitric oxide modulation of focal adhesions in endothelial cells. Am J Physiol 276, C1271–1281 (1999).

42. Takahashi, M. et al. Nitric oxide attenuates adhesion molecule expression in human endothelial cells. Cytokine 8, 817–821 (1996).

43. Kim, N.N., Villegas, S., Summerour, S.R. & Villarreal, F.J. Regulation of cardiac fibroblast extracellular matrix production by bradykinin and nitric oxide. J Mol Cell Cardiol 31, 457–466 (1999).

44. Hovater, M.B., Ying, W.Z., Agarwal, A. & Sanders, P.W. Nitric oxide and carbon monoxide antagonize TGF-beta through ligand-independent internalization of TbetaR1/ALK5. Am J Physiol Renal Physiol 307, F727–735 (2014).

45. Lee, S.W., Choi, H., Eun, S.Y., Fukuyama, S. & Croft, M. Nitric oxide modulates TGF-beta-directive signals to suppress Foxp3+ regulatory T cell differentiation and potentiate Th1 development. J Immunol 186, 6972–6980 (2011).

46. Kanno, Y., Into, T., Lowenstein, C.J. & Matsushita, K. Nitric oxide regulates vascular calcification by interfering with TGF-signalling. Cardiovasc Res 77, 221–230 (2008).

47. Zhou, X. & He, P. Improved measurements of intracellular nitric oxide in intact microvessels using 4,5-diaminofluorescein diacetate. Am J Physiol Heart Circ Physiol 301, H108–114 (2011).

48. Sulzmaier, F.J., Jean, C. & Schlaepfer, D.D. FAK in cancer: mechanistic findings and clinical applications. Nat Rev Cancer 14, 598–610 (2014).

49. Westhoff, M.A., Serrels, B., Fincham, V.J., Frame, M.C. & Carragher, N.O. SRC-mediated phosphorylation of focal adhesion kinase couples actin and adhesion dynamics to survival signaling. Mol Cell Biol 24, 8113–8133 (2004).

50. Mitra, S.K. & Schlaepfer, D.D. Integrin-regulated FAK-Src signaling in normal and cancer cells. Curr Opin Cell Biol 18, 516–523 (2006).

51. Bouchard, V. et al. Fak/Src signaling in human intestinal epithelial cell survival and anoikis: differentiation state-specific uncoupling with the PI3-K/Akt-1 and MEK/Erk pathways. J Cell Physiol 212, 717–728 (2007).

52. Nguyen, T.H. & Barr, F.G. Therapeutic Approaches Targeting PAX3-FOXO1 and Its Regulatory and Transcriptional Pathways in Rhabdomyosarcoma. Molecules 23 (2018).

53. Wachtel, M. & Schafer, B.W. PAX3-FOXO1: Zooming in on an “undruggable” target. Semin Cancer Biol 50, 115–123 (2018).

54. Taylor, J.G.t., et al. Identification of FGFR4-activating mutations in human rhabdomyosarcomas that promote metastasis in xenotransplanted models. J Clin Invest 119, 3395–3407 (2009).

55. Ginsberg, J.P., Davis, R.J., Bennicelli, J.L., Nauta, L.E. & Barr, F.G. Up-regulation of MET but not neural cell adhesion molecule expression by the PAX3-FKHR fusion protein in alveolar rhabdomyosarcoma. Cancer Res 58, 3542–3546 (1998).

56. Sunkel, B.D. et al. Evidence of pioneer factor activity of an oncogenic fusion transcription factor. iScience 24, 102867 (2021).

57. Lambert, A.W., Pattabiraman, D.R. & Weinberg, R.A. Emerging Biological Principles of Metastasis. Cell 168, 670–691 (2017).

58. Regina, C. et al. Negative correlation of single-cell PAX3:FOXO1 expression with tumorigenicity in rhabdomyosarcoma. Life Sci Alliance 4 (2021).

59. Han, S.J., Oak, Y., Groisman, A. & Danuser, G. Traction microscopy to identify force modulation in subresolution adhesions. Nature methods 12, 653 (2015).

60. Mittal, N. et al. Myosin-independent stiffness sensing by fibroblasts is regulated by the viscoelasticity of flowing actin. Communications Materials 5, 6 (2024).

61. Han, S.J., Oak, Y., Groisman, A. & Danuser, G. Traction microscopy to identify force modulation in subresolution adhesions. Nature methods 12, 653–656 (2015).

